# Long-range GABAergic projections contribute to cortical feedback control of sensory processing

**DOI:** 10.1101/2020.12.19.423599

**Authors:** Camille Mazo, Antoine Nissant, Soham Saha, Enzo Peroni, Pierre-Marie Lledo, Gabriel Lepousez

## Abstract

Cortical sensory areas send excitatory projections back to earlier stage of sensory processing. Here, we uncover for the first time the existence of a corticofugal inhibitory feedback between two sensory areas, paralleling the well-documented excitatory feedback. In the olfactory system, we reveal that a subpopulation of GABAergic neurons in the anterior olfactory nucleus and anterior piriform cortex target the olfactory bulb. These long-range inhibitory inputs synapse with both local and output olfactory bulb neurons, mitral and tufted cells. Optogenetic stimulation coupled to in vivo imaging and network modeling showed that activation these inhibitory inputs drives a net subtractive inhibition of both spontaneous and odor-evoked activity in local as well as mitral and tufted cells. Further, cortical GABAergic feedback stimulation enhanced separation of population odor responses in tufted cells, but not mitral cells. Targeted pharmacogenetic silencing of cortical GABAergic axon terminals in the OB impaired discrimination of similar odor mixtures. We propose here that cortical GABAergic feedback represents a new circuit motif in sensory systems, involved in refining sensory processing and perception.

## Introduction

Recent advances in genetic tools applied to cell- and circuit-tracing has allowed for the discovery of an increasing number of long-range GABAergic projection neurons in the cortex – where they may constitute 1-10% of the total GABAergic neurons in mice, rats, cats and monkeys (Higo, 2009; Higo et al., 2007; McDonald and Burkhalter, 1993; Tomioka and Rockland, 2007; Tomioka et al., 2005). Long-range projecting GABAergic neurons express a variety of classical markers for interneurons (Caputi et al., 2013; Melzer and Monyer, 2020; Urrutia-Piñones et al., 2022), sometimes forming intermingled populations within a single structure, where they exhibit distinct connectivity and exert various functions. For instance, bidirectional GABAergic projections between the hippocampus and entorhinal cortex synchronize the rhythmic network activity and gate spike-timing plasticity (Caputi et al., 2013; Melzer and Monyer, 2020), cortico-striatal and cortico-amygdala GABAergic projections regulate spike generation and excitability of their postsynaptic target and influence locomotion as well as reward coding (Lee et al., 2014; Melzer et al., 2017; Rock et al., 2016)

In mammalian sensory systems, external stimuli trigger a feedforward flow of information from the sensory organ to the primary and higher-order sensory cortices via a set of subcortical structures, thereby defining a hierarchy between sensory brain regions. In parallel, higher-order cortical sensory areas send top-down information to lower-order areas, constantly shaping early information processing. Such feedback is thought to convey contextual information and predictions to lower areas, not only playing a decisive role in selective attention and object expectation, but also in the encoding and recall of learned information (Gilbert and Li, 2013; Keller and Mrsic-flogel, 2018; Rao and Ballard, 1999). Top-down cortical feedback projections are thought to be exclusively mediated by glutamatergic neurons, while GABAergic neurons are in turn frequently referred to as exclusively mediating local information processing (Isaacson and Scanziani, 2011). In the present study, we challenge that view by investigating whether sensory cortical circuits can also parallelize excitatory and inhibitory top-down projections.

In the olfactory system, olfactory sensory neurons project to the olfactory bulb (OB), in the glomerular layer (GL) where they form synapses with apical dendrites of mitral and tufted cells (MCs and TCs, respectively), the output projection neurons of the OB. MC and TC activity is shaped by a large population of local GABAergic interneurons which synapse onto their apical or lateral dendrites. The anterior olfactory nucleus (AON) and the anterior piriform cortex (APC) — forming the anterior olfactory cortex (AOC) — is the primary recipient of OB outputs. Like the cortico-thalamic feedback pathway, the AOC send extensive glutamatergic projections back to the OB (Boyd et al., 2012; Carson, 1984; Davis and Macrides, 1981; Haberly and Price, 1978b, 1978a; Luskin and Price, 1983; Markopoulos et al., 2012; Mazo et al., 2017; de Olmos et al., 1978; Padmanabhan et al., 2016; Shipley and Adamek, 1984). Glutamatergic feedback projections from the AOC target virtually all types of neurons in the OB and induce robust disynaptic inhibition onto MCs and TCs (Boyd et al., 2012; Markopoulos et al., 2012; Mazo et al., 2016; Nissant et al., 2009; Strowbridge, 2009). These reciprocal connections between the OB network and the AOC are important for proper oscillations in the OB (Kay, 2014; Martin et al., 2006), decorrelation of OB output activity (Otazu et al., 2015), inter-hemispheric coordination (Grobman et al., 2018) and modulate odor perception threshold (Soria-Gómez et al., 2014) and odor-association learning (Wang et al., 2020) in a context-dependent manner (Wu et al., 2020). The OB additionally receives external inputs from neuromodulatory systems. Specifically, the basal forebrain sends GABAergic axons to the OB (Gracia-Llanes et al., 2010; Zaborszky et al., 1986) where they form synapses exclusively onto inhibitory neurons (Hanson et al., 2020; Sanz Diez et al., 2019; Villar et al., 2021). Optogenetic stimulation of basal forebrain GABAergic axons in the OB results in a bidirectional modulation of MCs, switching from an inhibitory to disinhibitory net effect in the presence of odor input (Böhm et al., 2020; Villar et al., 2021). Basal forebrain GABAergic projections have also been shown to influence OB oscillations, MC spike synchronization (Villar et al., 2021) and olfactory discrimination (Nunez-Parra et al., 2013).

Here we reveal that in addition to the cortical glutamatergic feedback, the AOC sends GABAergic projections back to the OB. Specifically, the AON *pars posterioralis* (AONp) form a particularly dense cluster of OB-projecting GABAergic neurons. Similar to their glutamatergic counterpart (Boyd et al., 2012; Markopoulos et al., 2012; Mazo et al., 2016) and in contrast to basal forebrain GABAergic inputs, we demonstrate that cortical GABAergic feedback forms synapses with MCs and TCs as well as deep-layer GABAergic interneurons, but spares GL GABAergic neurons. In awake mice, long-range GABAergic projection stimulation entrained beta oscillations in the OB. Cortical GABAergic feedback drives a net inhibition of both spontaneous and odor-evoked activity in local and output neurons, as predicted by network modeling. Further, cortical GABAergic feedback stimulation separated population odor responses in TCs, but not MCs. At the behavioral level, silencing of cortical GABAergic projections impaired fine odor discrimination of close binary mixture of enantiomers. Lastly, cortico-subcortical GABAergic projections are also observed between the primary somatosensory cortex (S1) and its respective lower-order thalamic nuclei.

## Results

### Anterior olfactory cortex sends GABAergic projections to the OB

To determine whether the AOC sends GABAergic projections back to the OB, in parallel to the well-described glutamatergic projections, we expressed different fluorescent reporters in the GABAergic and glutamatergic populations of the AOC. Using transgenic mice expressing the Cre recombinase under the vesicular GABA transporter VGAT (VGAT-Cre), we employed a conditional genetic approach to restrict expression of eYFP in GABAergic neurons while expressing mCherry in excitatory neurons using the CaMKIIa promoter (Fig. 1a). Both GABAergic and glutamatergic axons were found in the OB but showed different innervation profiles. While GABAergic axons accumulated preferentially in the superficial granule cell layer (GCL) to the mitral cell layer (MCL), glutamatergic axons were more concentrated in the deep GCL and their density progressively decreased towards the MCL (Fig 1b). To control for the specific expression of the conditional vectors, we injected the same virus mix in the AOC of wild-type mice and found no expression of the conditional fluorophore eYFP in the AOC (Supplemental Fig 1a). Further, to confirm the GABAergic nature of the labeled cortical axons in the OB, we injected a conditional virus expressing GFP and Synaptophysin fused with mRuby in the AOC of VGAT-Cre mice. In the OB, synaptohysin-mRuby^+^ presynaptic boutons colocalized extensively with VGAT immunostaining (Fig 1c).

**Figure 1.**
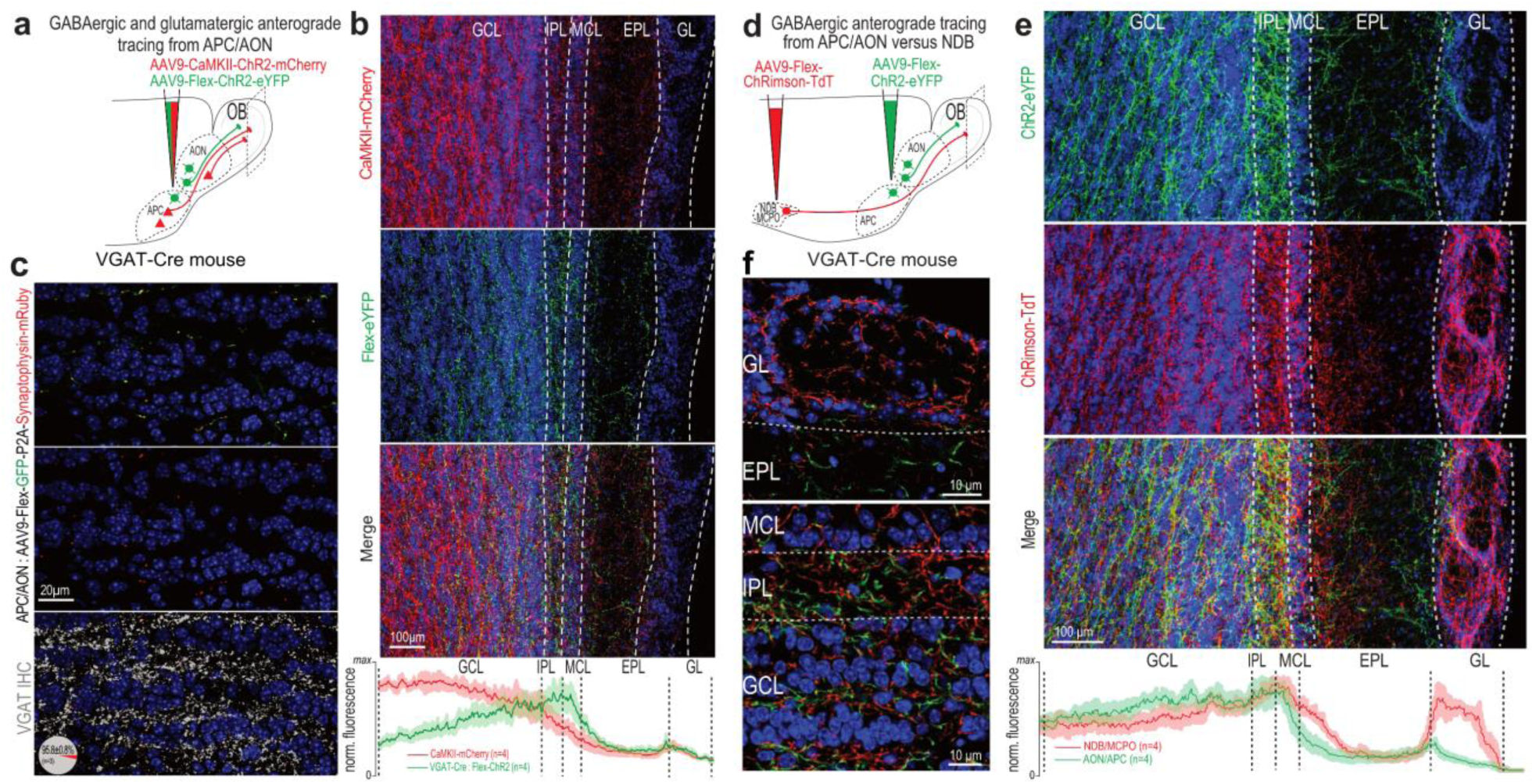
The olfactory cortex sends GABAergic projections back to the OB. **a**, Viral strategy for comparative anterograde labeling of glutamatergic (CaMKII-ChR2-mCherry) and GABAergic (Flex-ChR2- eYFP) axons in the OB from the AON/APC in a VGAT-Cre mice. **b,** Confocal images exhibiting the laminar profile of OB innervation by glutamatergic (top, red) or GABAergic (middle, green) axons from the AON/APC. Bottom, merge. Bottom plot, normalized axon fluorescence intensity from glutamatergic (red) versus GABAergic (green) across OB layers. Mean (solid line) ± sem (shaded). n = 4 mice. **c,** High magnification of the GCL of VGAT-Cre mice injected with AAV9-Flex-GFP-P2A-Synaptophysin-mRuby in the AON/APC. Top, GABAergic axon shafts (green) and their presynaptic mRuby-positive boutons (red). Middle, Synaptophysin-mRuby channel only. Bottom, immunohistochemistry (IHC) against VGAT is overlayed with the Synaptophysin-mRuby signal. 98 ± 0.8 % of the putative boutons colocalize with VGAT staining (n = 3 mice). **d**, Comparative anterograde labeling of AON/APC (ChR2-eYFP) versus NDB/MCPO (ChRimson-TdTomato) GABAergic axons in the OB using Cre-dependent AAV injection in VGAT-Cre mice. **e**, Confocal images exhibiting the laminar profile of OB innervation by GABAergic axons from the AON/APC (top, green) or NDB/MCPO (middle, red). Bottom, merge. Bottom plot, normalized axon fluorescence intensity from AON/APC (green) versus NDB/MCPO (red) across OB layers. Mean (solid line) ± sem (shaded). n = 4 mice. **f**, Higher magnification of **e** in the different OB layers. NDB/MCPO and AON/APC GABAergic axons are intermingled but distinct in the OB. Blue, DAPI. GL, glomerular Layer; EPL, external plexiform layer; MCL, mitral cell layer; IPL, internal plexiform layer; GCL, granule cell layer.

Another identified source of GABAergic inputs to the OB originates from the nucleus of the diagonal band and magnocellular preoptic area (NDB/MCPO)(Gracia-Llanes et al., 2010; Kunze et al., 1992a, 1992b; Zaborszky et al., 1986). We directly compared the OB innervation patterns of GABAergic axons from the olfactory cortex vs. NDB/MCPO using conditional virus expression in both brain regions (Fig. 1d). NDB/MCPO and AOC axon innervation patterns were strikingly different in the GL (Fig. 1e,f). While NDB/MCPO profusely innervated the GL (Böhm et al., 2020; Hanson et al., 2020; Nunez-Parra et al., 2013; Sanz Diez et al., 2019; Villar et al., 2021), AON/APC projections were restricted to the internal part of the GL. NDB/MCPO axons also appeared to innervate more the inner part of the external plexiform layer (Böhm et al., 2020) (EPL).

Taken together, these results show that the AOC sends GABAergic axons back to the OB, and these projections are distinguishable from the well-established cortico-bulbar glutamatergic projections and from the basal forebrain GABAergic projections.

### Identity of the long-range GABAergic projections to the OB

To identify the source(s) of cortical GABAergic feedback to the OB, we employed a conditional retrograde labeling approach. A Herpes Simplex Virus (HSV) expressing GCaMP6f in a Cre-dependent manner was injected in the OB of VGAT-Cre mice (Fig. 2a). Retrogradely-labeled cells were found mainly in the AON, APC and NDB/MCPO, and occasionally in the posterior piriform cortex (PPC) and tenia tecta (TT), but not in the olfactory tubercle (OT) ― a large striatal GABAergic structure of the olfactory system (Fig. 2a-c). Following unilateral injection, retrogradely labeled cells were found only in the ipsi-lateral, but not contra-lateral side of the injection (Supplemental Fig. 2a). Retrogradely-labeled cells were not uniformly distributed within the AOC. A large proportion of the labeled cells were concentrated in the AON *pars posterioralis* (AONp) — the most caudal part of the AON, located in between the APC and the OT (Fig. 2c). An appreciable fiber tract was often observed between the AONp and the NDB/MCPO (Fig. 2a), yet the newly identified cluster of GABAergic projection neurons was not a rostral extension of striatal or pallidal territory as it was not intermingled with neurons expressing the acetylcholine-synthesizing enzyme ChAT – in contrast to OB-projecting GABAergic neurons of the NDB/MCPO (Hanson et al., 2020; Villar et al., 2021)(Supplemental Fig. 2b). To confirm these observations based on retrograde viral vectors, we combined conventional retrograde labeling of OB-projecting neurons using cholera toxin subunit-B conjugated to a red fluorophore (CTB) with somatic viral labeling of GABAergic neurons of the AON/APC. Likewise, dually-labeled cells were found scattered in the AOC, with a higher density in the AONp (Supplemental Fig. 2c).

**Figure 2.**
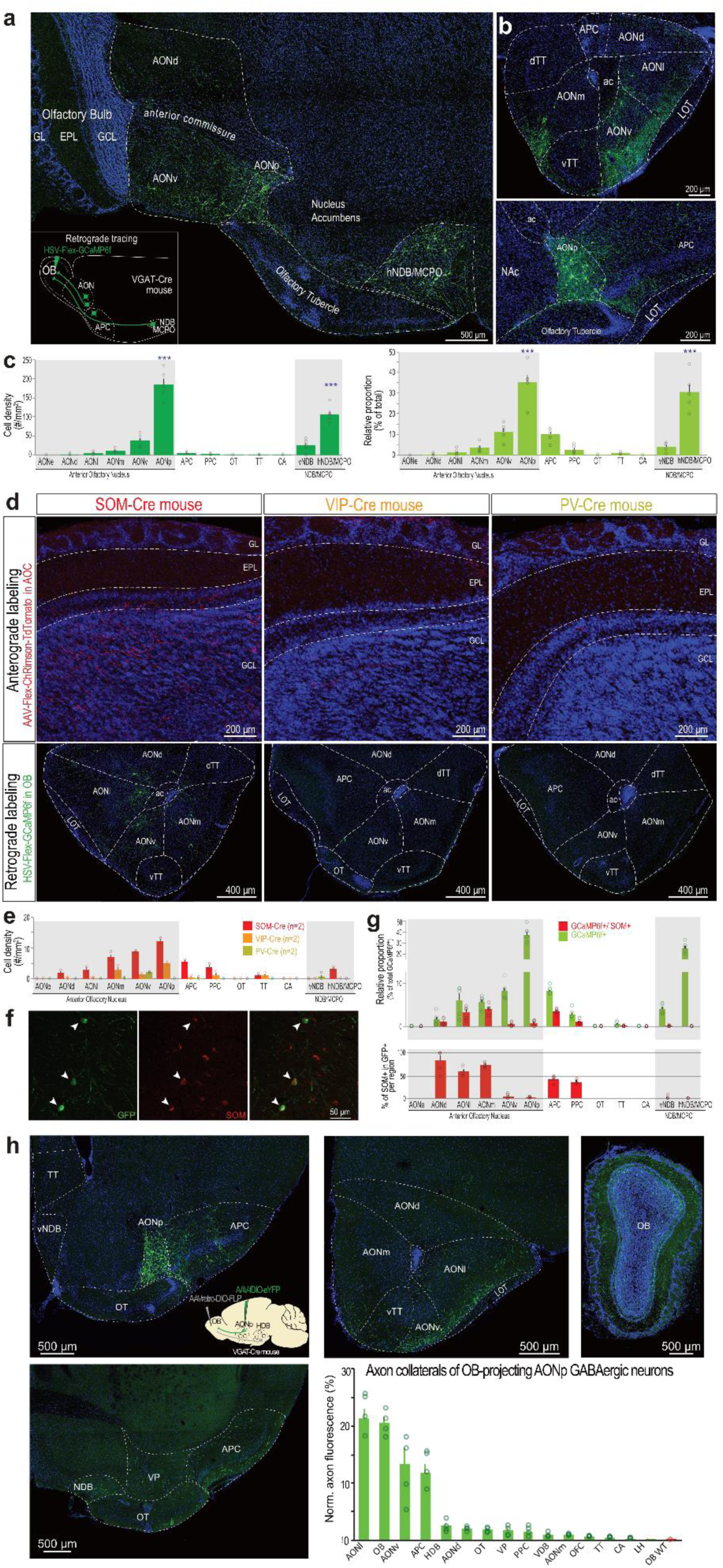
Anatomical and neurochemical identification of the OB-projecting GABAergic cells in the anterior olfactory cortex. **a**, OB-projecting GABAergic cells in a sagittal slice. Inset: schematic of the injection. **b**, Coronal slices through the AON (top) and APC/AONp (bottom). **c,** Cell density (left) and relative proportion (right) of the OB-projecting GABAergic cells (**n = 2134 cells from 5 mice**). AONp and hNDB/MCPO contain the largest density and proportion of cells (density: one-way ANOVA, F(12,52) = 75.45, p = 10^-28^ Tukey multiple comparison test, AON vs each individual areas, p < 10^-7^, hNDB/CMPO vs all the other individual areas, p < 10^-7^; proportions: one-way ANOVA, F(12,52) = 40.86, p = 10^-22^, Tukey multiple comparison test, AON vs hNDB/MCPO, p = 0.86, AON vs all the other individual areas, p < 10^-7^, hNDB/CMPO vs all the other individual areas, p < 10^-7^). **d**, Top, anterograde labeling of SOM-, VIP- and PV-axons in the OB (ChRimson-tdTomato). Sagittal sections through the OB. Bottom, retrograde labeling of OB-projecting cells (GCaMP6f) in SOM-, VIP- and PV-Cre mice. Coronal sections through the AON. **e**, Cell density of the retrogradely-labeled cells as in **c** but for SOM-Cre (red; n = 2 mice), VIP-Cre (orange; n = 2 mice) and PV-Cre mice (green; n = 2 mice). **f**, Example SOM labeling of OB-projecting GABAergic neurons in the AON. Arrowheads point to co-labeled cells. **g,** Top, quantification of co-labeled cells across cortical olfactory regions (n = 1075 cells from n = 5 mice). Bottom, percentage of SOM^+^ cells among all the GCaMP^+^ cells in that region. **h**, Selective expression of eYFP in OB-projecting GABAergic neurons of the AOC was achieved using a double-conditional strategy. In VGAT-Cre mice, a retrograde AAV conditionally expressing the flipase recombinase (FLP) was injected in the OB (AAVretro-DIO-Flp) and a flipase-dependent AAV expressing eYFP was injected in the AONp (AAV-fDIO-ChR2-eYFP), restricting somatic labeling mainly to the AONp (85.1 ± 2.5% of labeled neurons in the AONp, 9.9 ± 1.4 in the APC, 3.2 ± 0.4% in AONv and 1.6±0.6 in AONl, n = 4). As a control, the AAV-fDIO-ChR2-eYFP was also injected in the NAc, a region not projecting to the OB. Axons were found in various anterior olfactory cortical areas (quantified in the lower right panel; n = 4 mice; OB WT is the specificity control described in Supplemental Fig. 3d). In **c, e, g**, and **h,** data is presented as mean ± sem across mice. Circles are individual mice. AONl, AON lateralis; AONd, AON dorsalis; AONv, AON ventralis; AONp, AON posterioralis; AONm, AON medialis; OT, olfactory tubercle; dTT, dorsal Tenia Tecta; vTT, ventral Tenia Tecta; LOT, lateral olfactory tract; hNDB/MCPO, horizontal limb of the nucleus of the diagonal band / magnocellular preoptic nucleus; vNDB, vertical limb of the nucleus of the diagonal band; CA, cortical amygdala; NAc, nucleus accumbens; ac, anterior commissure.

We found both spiny and aspiny neurons in the cortical OB-projecting GABAergic neurons (Supplemental Fig. 3a). We thus set out to substantiate these observations and precise their neurochemical nature. Somatostatin (SOM), parvalbumin (PV) and the vasoactive intestinal peptide (VIP) characterize the vast majority of GABAergic neurons in the cortex and have been reported in largely non-overlapping populations in the AON (Kay and Brunjes, 2014) and APC (Suzuki and Bekkers, 2010). To identify the marker preferentially expressed by OB-projecting cortical GABAergic neurons, we first injected an AAV-Flex-ChR2-TdTomato in the AON/APC of SOM-Cre, VIP-Cre or PV-Cre mice. Substantial axonal innervation in the OB was observed in SOM-Cre mice, while very sparse fibers were detected in PV-Cre and VIP-Cre mice (Fig. 2d). Reciprocally, injection of a Cre-dependent retrograde vector in the OB led to a denser number of neurons in SOM-Cre than in VIP-Cre or PV-Cre mice (Fig. 2d,e). However, we did not observe a predominant labeling in the AONp, in contrast to results obtained in VGAT-Cre mice. To confirm this observation, we performed immunostaining against the protein SOM in retrogradely-labeled GABAergic neurons (VGAT-Cre mice; Fig. 2f). In the olfactory cortex, we found that a substantial fraction of OB-projecting GABAergic neurons was co-labeled with SOM in the AONd, AONl, AONm, APC and PPC, but not in the AONv or AONp (Fig 2g). In SOM-Cre mice, 95.5 ± 0.5% of the genetically-labeled neurons were dually-labeled with SOM immunostaining, confirming the efficiency of our SOM immunohistological labeling (Supplemental Fig. 1b). Focusing on the AONp, the densest source of projection neurons, we performed further immunohistological characterization for GABAergic neuron markers and found that calbindin colocalized more than any other marker tested, yet still to a modest degree (Supplemental Fig. 3b,c).

We also wonder whether OB-projecting GABAergic neurons send axon collaterals elsewhere in the brain. Using a double-conditional strategy based on FLP and Cre recombinase, we specifically restricted the expression of eYFP to GABAergic OB-projecting neurons of the AONp (Fig 2h, Supplemental Fig. 3d). In addition to the OB, these AONp GABAergic neurons innervate the AONl, AONv and APC, but no sizeable axonal arborization was found in the NDB/MCPO or lateral hypothalamus. In conclusion, inhibitory projection neurons were found scattered in the AOC, with a substantial cluster located in the AONp. These GABAergic neurons innervate preferentially the olfactory system and have limited projections in non-olfactory areas.

### The primary somatosensory cortex also sends GABAergic projections to the somatosensory thalamus

We wondered whether such inhibitory cortical feedback motifs down the hierarchy existed in other sensory systems. In the primary somatosensory cortex (S1), we performed a similar dual anterograde labeling of deep-layers GABAergic and glutamatergic neurons (Supplemental Fig. 4a). GABAergic axons were found alongside glutamatergic axons in the lower-order (ventroposterior medial and lateral, VPM and VPL) and higher-order (posteriomedial, POm) somatosensory thalamic nuclei (Supplemental Fig. 4b). GABAergic cortico-thalamic projections intermingled with glutamatergic projections and did not seem to project to other thalamic territories. These results indicate that corticofugal inhibitory projections might be a more common motif than previously thought in sensory pathways.

### Cortical GABAergic projections target both OB principal cells and interneurons

We next tested whether cortical GABAergic inputs form functional synapses onto OB neurons and whether GABAergic inputs exhibit target selectivity. Channelrhodopsin-2 (ChR2) was expressed selectively in the GABAergic cells of the AON and APC — the 2 regions consisting of ∼95% of the OB-projecting cortical neurons (Figure 2c) — and whole-cell recordings were obtained in acute OB slices (Fig 3a). In responsive neurons (74/177), light stimulation of GABAergic axons evoked short-latency post-synaptic currents (PSCs; Supplemental Table 1), consistent with monosynaptic events. PSCs were characterized by a current-voltage linear relationship reversing at the reversal potential for chloride (∼-75mV; Fig 3b). and were unchanged in presence of the AMPA receptor antagonist NBQX (10 µM) but were completely abolished by the GABA_A_ receptor antagonist gabazine (SR95531, 10 µM; Fig 3c,d), confirming the GABAergic nature of the PSCs.

**Figure 3.**
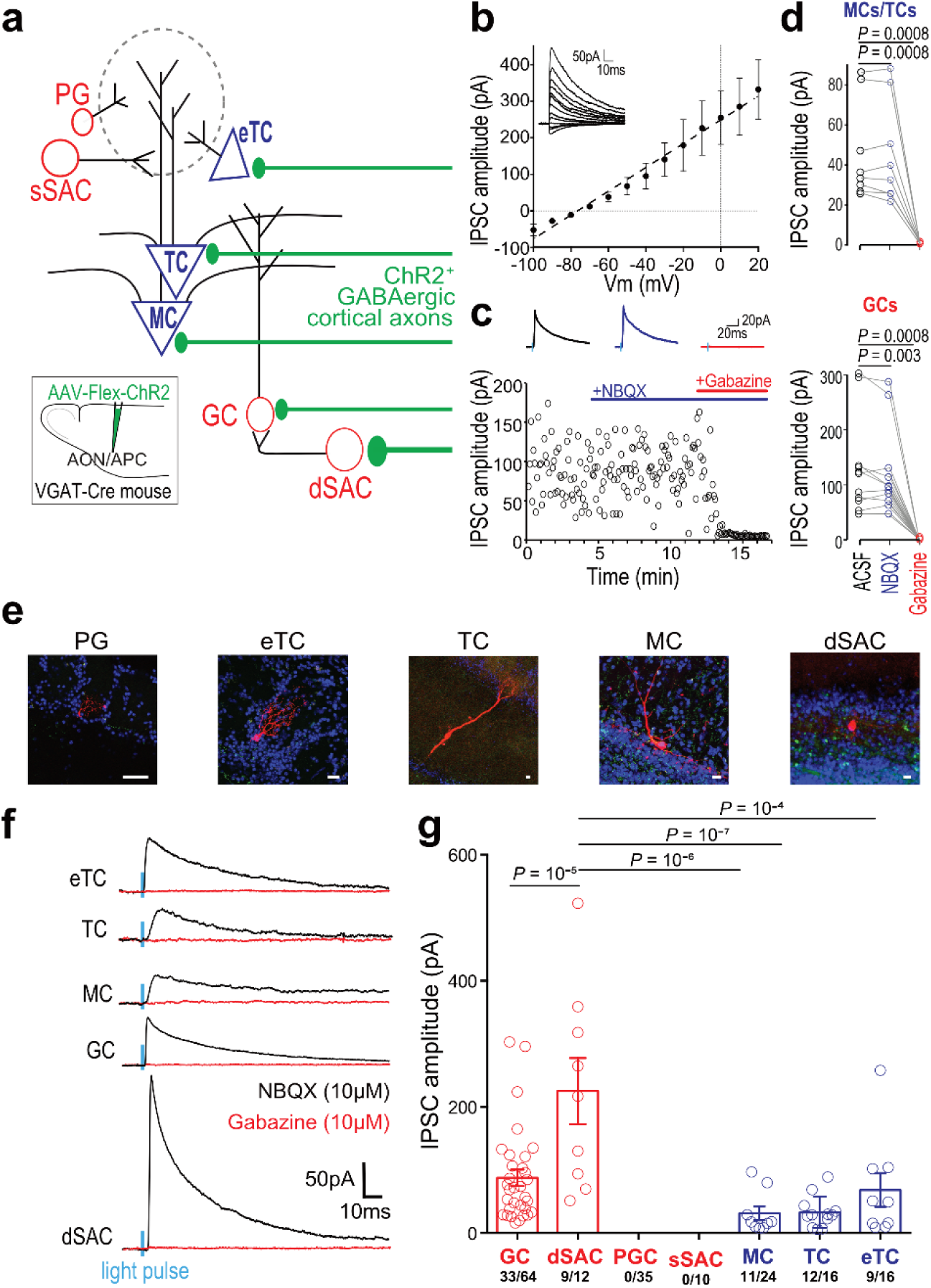
Cortico-bulbar GABAergic axons form functional synapses with inhibitory and excitatory neurons in the OB. **a**, Recording schematic. Periglomerular (PG) cells, external tufted cells (eTCs), superficial short-axon cells (sSAC), tufted cells (TCs), mitral cells (MCs), granule cells (GCs) and deep short-axon cells (dSACs) were patched and GABAergic feedback axons expressing ChR2 (inset) were light-stimulated (2 ms). Width of the axon shafts indicates connection probability **b**, PSCs recorded in GCs (n = 4) at different holding potentials, in the presence of NBQX (10 µm). Responses reversed at ∼-75 mV, consistent with GABAergic receptor activation. In the rest of the figure, V_c_ = 0 mV. **c,** IPSC amplitudes in an example GC in the presence of ACSF, NBQX or gabazine. **d,** PSCs were resistant to NBQX application (10 µM) but completely abolished by gabazine (SR95531, 10 µM) in MCs/TCs (top) and GCs (bottom; GCs: One-way ANOVA, F(2,33) = 9.82, p = 10^-4,^ with Tukey’s post-hoc test, n = 11; MCs/TCs: One-way ANOVA, F(2,21) = 12.56, p = 0.0003 with Tukey’s post-hoc test, n = 8). **e**, Representative images of the patched neurons. Neurons were first visualized and identified online. GCs and sSACs morphology were not reconstructed *post hoc* because pipette withdrawal after recording did not preserve the integrity of the cells. Scale bars are 10 µm. **f,** Representative trial-averaged IPSCs in cells recorded at V_c_ = 0 mV. Responses were systematically blocked in gabazine. Blue, light-pulse; black, recordings in NBQX (10 µM); red, recordings in gabazine (SR95531, 10 µM). **g**, IPSC amplitudes across the cell tested (One-way ANOVA, F(4,68) = 11.23, p = 10^-7^, with Tukey’s post-hoc test). Blue bars, excitatory neurons; Red bars, inhibitory neurons; circle, individual cell. Data presented as mean ± sem.

We next investigated the target specificity of these cortical GABAergic inputs. OB neurons were classified according to their intrinsic properties, morphology, soma size, and laminar position in OB slice (Fig 3e; see Material and Methods). Inhibitory PSCs (IPSCs) were detected in roughly half of the excitatory neurons tested (MCs, TCs, and eTCs). Among GCL inhibitory neurons, we found a stronger connectivity and larger current amplitude in dSACs compared to GCs (Fig. 3f,g), reminiscent of previous observations with glutamatergic feedback (Boyd et al., 2012; Markopoulos et al., 2012). In contrast, glomerular inhibitory neurons were spared, consistent with the lower density of axons observed in the glomerular region (Fig. 1b, e). To confirm a direct synaptic connection from cortical GABAergic neurons, we performed rabies-based retrograde monosynaptic tracing from MCs and TCs (Wickersham et al., 2007) (Supplemental Fig. 5a). Monosynaptic retrogradely labeled neurons were observed in the AOC and some colocalized with the GABA synthesizing enzymes glutamic acid decarboxylase 67 (GAD67; Supplemental Fig. 5a). We also found monosynaptic retrogradely labeled GABAergic cells in the AOC with GCL GABAergic interneurons as the starter cell population (Supplemental Fig. 5b). Thus, cortical GABAergic feedback provides direct functional inputs to a variety of OB neurons, both inhibitory and excitatory.

### Cortical GABAergic inputs influence OB network oscillations

Long-range cortical GABAergic neurons have been repeatedly proposed to play a role in tuning network oscillations and synchronizing distant brain areas (Melzer and Monyer, 2020). To explore the functional role of cortical GABAergic projections to the OB, we first investigated to what extent this cortico-bulbar GABAergic pathway can influence oscillatory regimes in the OB. Network oscillations are prominent in the OB and can be subdivided into different frequency bands, theta (1-12 Hz) beta (15-40 Hz) and gamma oscillations (40-100 Hz). We coupled repetitive optogenetic stimulation of cortical GABAergic axons at different frequencies with local field potential (LFP) recordings in the OB of awake VGAT-Cre mice to investigate if the OB network respond maximally, or resonate, at specific driving frequencies (Supplemental Fig. 6a). Spontaneous beta oscillations, but not theta or gamma, were amplified upon light stimulation of GABAergic cortical axons. Specifically, light stimulation at 33 Hz increased beta band frequencies, while stimulation at 10 or 66 Hz had no significant effect (Supplemental Fig. 6b,c). Thus, long-range cortical GABAergic projections tune OB oscillations and specifically enhance beta oscillations when entrained at beta frequencies.

### Optogenetic activation of cortical GABAergic inputs inhibits OB GCL interneurons *in vivo*

To assess the functional impact of the cortical GABAergic feedback on its main target layer (GCL), we employed fiber photometry in freely moving mice. The volume fluorescence of GCaMP6f-expressing GCL GABAergic neurons was continuously recorded using an optic fiber implanted above the GCaMP6f injection site, while AOC GABAergic projections were light-stimulated in the ventral OB (Fig. 4a). Using the red-shifted opsin ChRimson, we could independently control GABAergic axons and avoid cross-excitation of GCaMP6f (Klapoetke et al., 2014; Mazo et al., 2016). ChRimson light stimulation at 10 Hz, 33 Hz or with a continuous light step (CL) produced a global reduction of spontaneous activity in GCL GABAergic neurons while red light stimulation *per se* did not alter spontaneous activity in control animals (expressing GCaMP6f in the GCL, but not ChRimson in GABAergic feedback; Fig. 4b). Increasing light stimulation frequency up to a continuous light pulse induced increasing inhibition of GCL neuron activity (Fig. 4b). This feature was still observed 1 s after light stimulation offset (Supplementary Fig. 7a).

**Figure 4.**
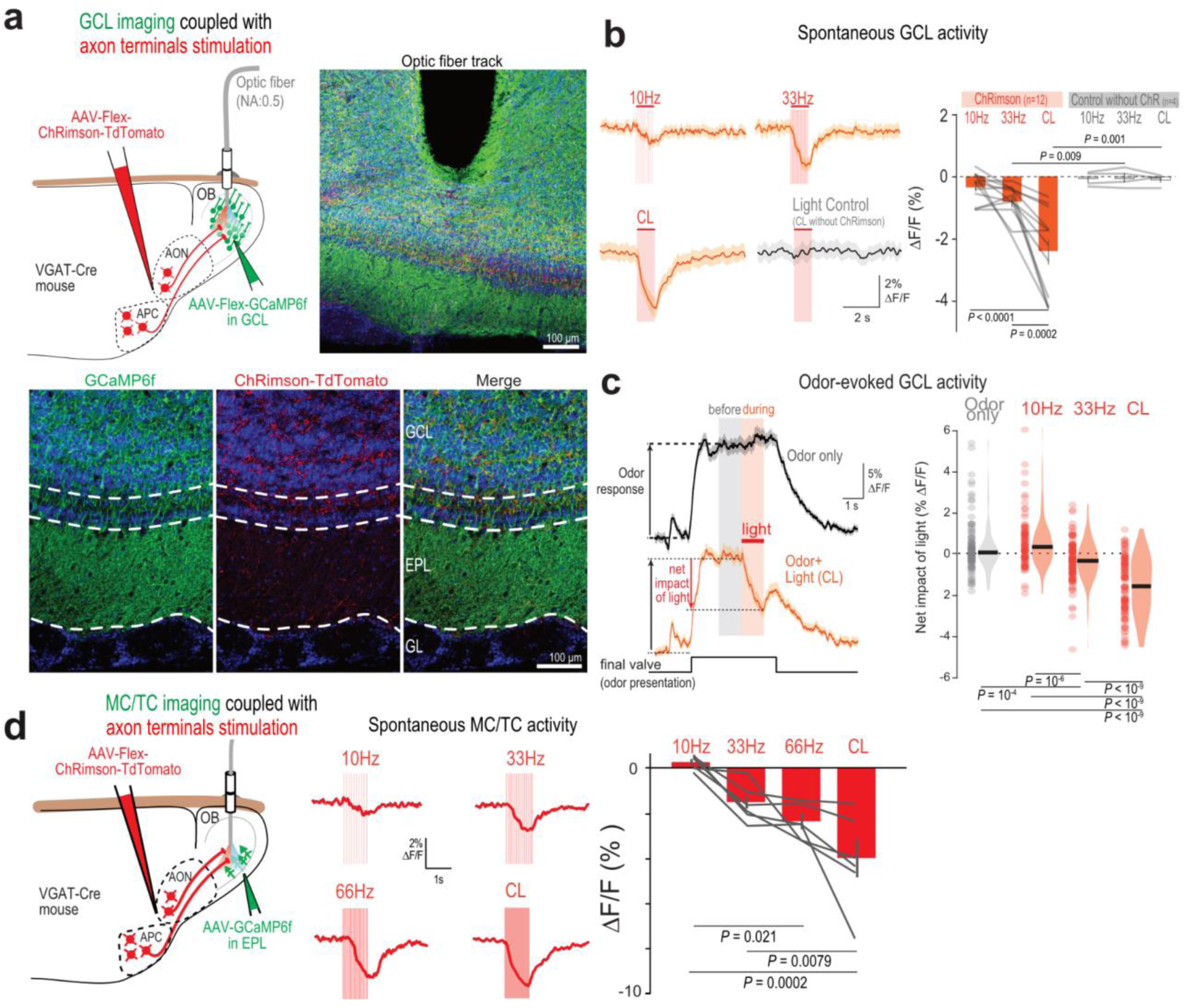
Cortico-bulbar GABAergic axons induce net inhibition onto GCL neurons and MC/TC populations *in vivo*. **a**, Cortical GABAergic axons were stimulated in the OB of freely moving mice. Right, Optic fiber for cortical GABAergic axon stimulation and interneurons calcium imaging in the ventral GCL. Bottom, GCaMP6f expression across OB layers and ChRimson in AON/APC GABAergic axons. Blue, DAPI. **b**, Light stimulation (1 s) of cortical GABAergic axons decreased GCL interneuron spontaneous population activity. Left, example trial-averaged responses to different light stimulation patterns. “Light control” is light illumination (CL) in mice lacking ChRimson expression. Right, Stimulation at 33Hz and CL, but not at 10 Hz, induced a significant change in mean fluorescence (RM-One-way-ANOVA with Tukey’s post-hoc test, F(2,22)=21.28, P<0.0001). Data presented as mean ± sem; gray lines, individual mice. **c**, Light stimulation (1 s) of cortical GABAergic axons decreased GCL interneuron odor-evoked population activity. Left, fluorescence signals during odor presentation only (black) and odor presentation coupled with light stimulation of cortical GABAergic axons (orange). Traces are trial-averaged example responses. Mean ± sem are represented. Right, Net impact of light on odor responses across light stimulation protocols (10 Hz, 33 Hz, CL) compared to no light stimulation (“odor only”), during light stimulation (left, orange). The net impact of light was measured as the difference between the mean odor-evoked response in the 1 s window before versus during light stimulation. Population odor response where inhibited by light stimulation of cortical GABAergic axons and inhibition magnitudes correlated with the stimulation strength during light stimulation (One-way-ANOVA with Tukey’s post-hoc test, F(3,416) = 62.98, p = 10^-3^). Violin plots are ks density estimates; black bar is median; circle, individual odor-recording site pair. **d,** MC/TC population responses to cortical GABAergic axon stimulation in the OB of freely moving mice utilizing fiber photometry (left). Middle: Representative trial-averaged traces of light-evoked inhibitory responses in MC/TC using different stimulation patterns. Right: Light stimulation of cortical GABAergic axons produced increasing inhibition with increasing stimulation frequency (RM-One-way ANOVA with Tukey’s post-hoc test, F(3,15)=12.20, P=0.0003, n = 6 recording sites in 4 mice). Data presented as mean ± sem; Gray, individual recording sites.

We next investigated the impact of GABAergic feedback stimulation on odor-evoked activity in the GCL. Odor stimulation induced a strong population response in GCL neurons (Wang et al., 2020) (Fig. 4c). We quantified the net decrease in Ca^2+^ activity relative to the period before light stimulation, within the same odor response (Fig. 4c). When compared with odor response dynamics without light stimulation (“odor only”), GABAergic feedback light stimulation effectively dampened odor responses with 33 Hz and CL, but not 10 Hz, stimulation patterns (Fig. 4c). CL light inhibition of odor responses outlasted the light stimulation period: 1 s after light stimulation offset, CL still caused a sustained inhibition of the odor-evoked activity (Supplemental Fig. 7a). Thus, cortical GABAergic axon stimulation efficiently drives inhibition of both spontaneous and odor-evoked activity in GCL GABAergic neurons.

### Cortical GABAergic inhibition to OB output neurons scales with the frequency of stimulation

Given that cortical GABAergic feedback synapses both on GCs and OB principal neurons, we wondered how GABAergic axon stimulation impacts MC/TC activity in vivo. We targeted our fiber photometry recordings to MCs/TCs and found that light stimulation of cortical GABAergic axons inhibited MC/TC activity (Fig. 4d). Increasing light stimulation frequency up to a continuous light pulse induced increasing inhibition of MC/TC activity (Fig 4d). In addition to GABAergic feedback-mediated inhibition, the AOC inhibits MCs and TCs through a disynaptic pathway: glutamatergic feedback drives GCs and PG cells which in turn inhibit MCs and TCs (Boyd et al., 2012; Markopoulos et al., 2012; Mazo et al., 2016) (Supplemental Fig. 7b,c). We thus wished to compare the differential frequency recruitment of inhibition resulting from stimulation of the monosynaptic cortical GABAergic vs. disynaptic cortical glutamatergic pathway. While GABAergic projection stimulation drove increasing MC/TC inhibition with increasing stimulation frequency, the MC/TC inhibition evoked by light stimulation of glutamatergic projection peaked at 33 Hz, implementing a bell-shaped, low-pass filtering of the excitatory drive (Supplemental Fig. 7c).

### Computational modeling of cortical inhibitory feedback on OB network

Optogenetic stimulation of GABAergic feedback axons results in a net inhibition of the two recurrently connected excitatory (MCs and TCs) and inhibitory neuron populations (GCs). To tackle this apparent paradoxical effect and further explore the outcomes of GABAergic feedback on OB neurons, we built a population model of the OB based on our experimental results. Our model consisted of reciprocally connected excitatory (MCs/TCs) and inhibitory subnetworks (GCs). GCs additionally receive inhibitory inputs from an additional inhibitory population (dSACs; Fig 5a). We computed the steady state of the network without and with GABAergic feedback over a range of GC-MC/TC synapse and feedback strength. The relative strength of the feedback on MCs/TCs and GCs was derived from our slice recording data (Fig 3). This parsimonious model showed that GABAergic feedback resulted in a net inhibition on both MCs/TCs and GCs (Fig 5c,d). Importantly, this effect was observed for a range of values consistent with GC and MC/TC firing rate observed in vivo (Fig 5b). We additionally observed that increasing GABAergic feedback stimulation strength produces stronger inhibition on MCs/TC and GCs, which mirrors our in vivo data with increasing stimulation frequency (Fig 4).

**Figure 5.**
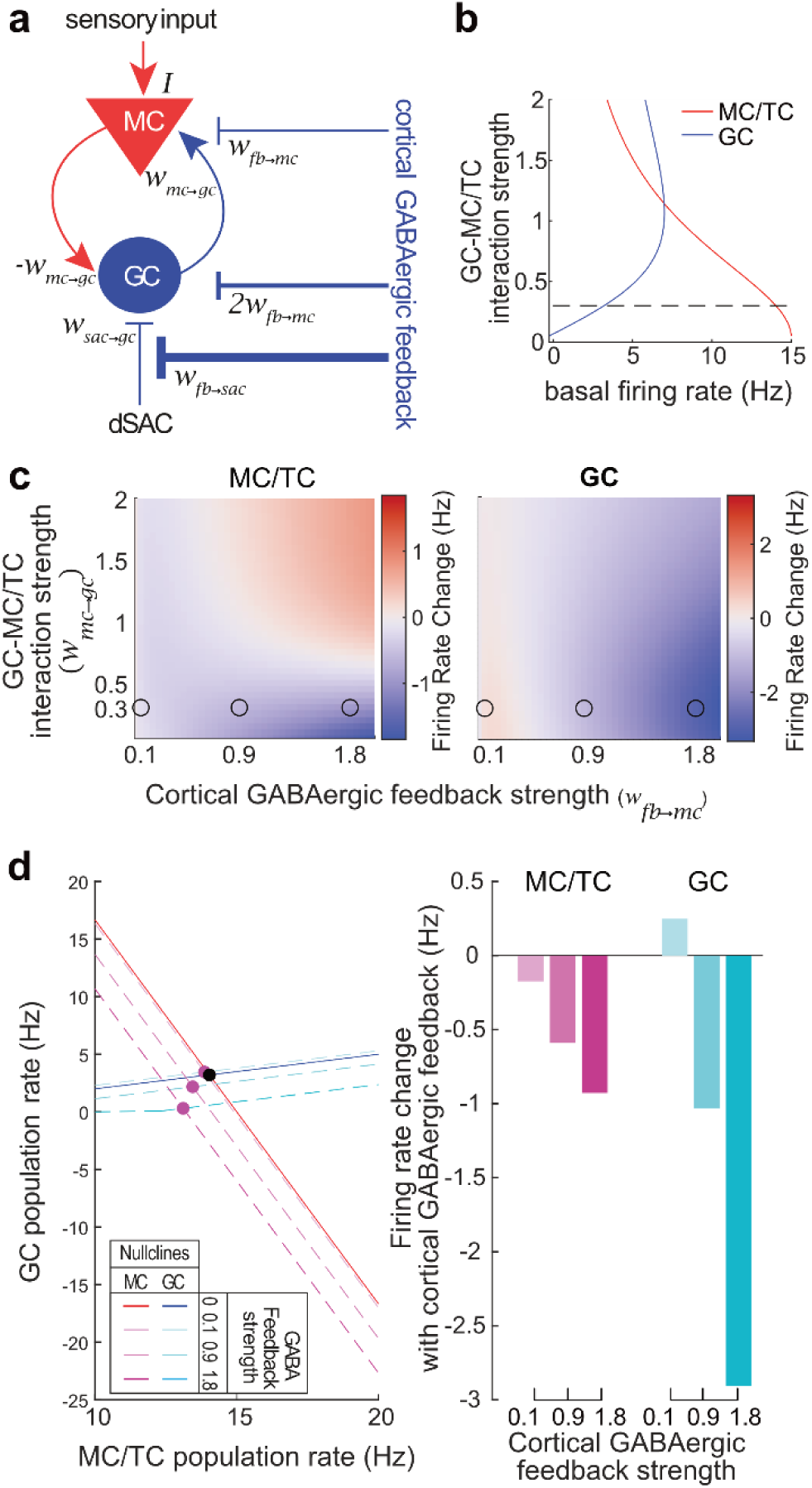
A parsimonious population model recapitulated the inhibition of both MC/TC and GC populations upon cortical GABAergic feedback stimulation. **a**, Schematic of a population model with excitatory (MCs/TCs, red) and inhibitory (GCs, blue) subnetworks. MC/TC population additionally receives external excitatory inputs (odor stimulation) and GC population inhibitory inputs from dSACs. Cortical GABAergic feedback directly inhibits MC/TC and GC populations and is twice stronger on GCs. It also shunts the inhibitory input from the dSAC to GC population. **b**, Basal firing rate of MC/TC and GC populations when the system is at the equilibrium (fixed point). Dashed line is the value *w_mc→gc_* chosen in d. **c**, Change in the firing rate upon cortical GABAergic feedback stimulation for a range of GC-MC interaction and feedback strength. Dots are the strength of the MC/TC-GC and GABAergic feedback used in d. **d,** Fixed point analysis of the system without and with different strength of cortical GABAergic feedback stimulation. Left, steady state in MC/TC and GC population is represented by the nullclines, (MC, solid red line without stimulation, dashed lines in shades of purple with feedback stimulation; GC, solid blue line without stimulation, dashed lines in shades of blue with feedback stimulation). Equilibrium of the system is obtained by the crossing of the nullclines (fixed point: without stimulation, black; with stimulation, purple). GC firing rate is rectified to avoid negative firing rates. Right, quantification of the change in firing rate for the MC/TC and GC populations for the different strength of the GABAergic feedback stimulation.

### Activation of cortical GABAergic inputs enhances the distance in TC population odor responses

Our fiber photometry data identified that the MC/TC population, taken as a whole, is inhibited upon light stimulation of cortical GABAergic axons. Are MCs and TCs similarly inhibited by GABAergic feedback? To resolve individual MCs and TCs, we performed two-photon Ca^2+^ recordings in awake, head-fixed mice. Based on our fiber photometry data, we used 33 Hz and CL stimulation to probe optimal regimes of cortical inhibitory drive. We therefore switched to the opsin ChIEF because it yields stronger currents upon long light pulses and a more naturalistic drive of GABAergic axons (Fig 6a) (Lin et al., 2009). The axon terminals were light-stimulated through the microscope’s objective, while the photomultiplier tube (PMT) shutter was closed and reopened 50 ms before and after light onset and offset. Due to the slow kinetics of GCaMP6s, we could capture Ca^2+^ events following the light offset and reopening of the shutter (Fig. 6c). MCs and TCs were identified based on the recording depth and the cytoarchitecture of each OB layer (Adam et al., 2014; Kikuta et al., 2013; Otazu et al., 2015; Sailor et al., 2016; Yamada et al., 2017).

**Figure 6.**
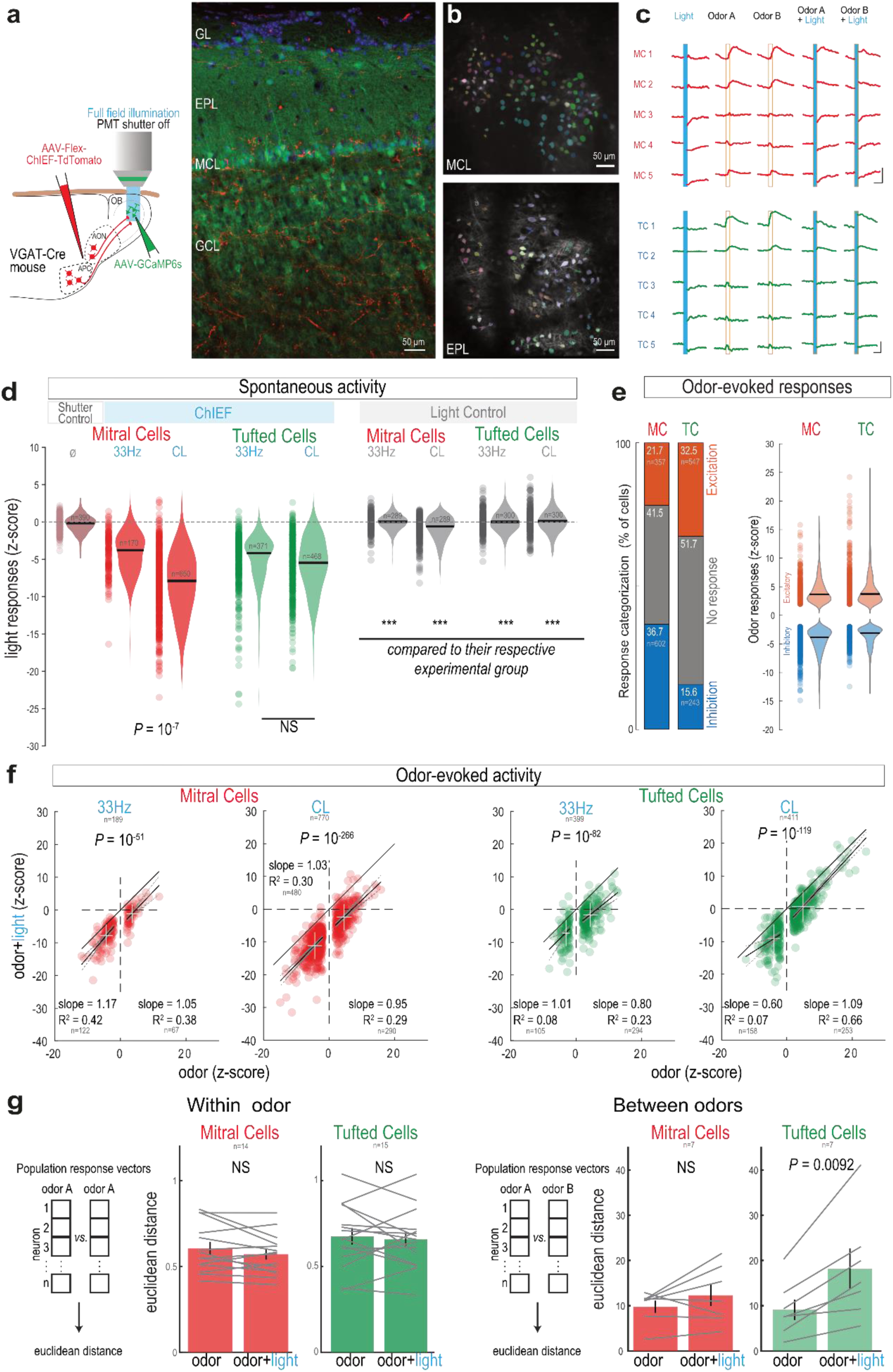
GABAergic feedback axon stimulation inhibits TC and MC activity *in vivo*. **a**, Two-photon imaging of TC and MC in awake mice coupled to full-field optogenetic stimulation of GABAergic cortical axons through the microscope objective. Right, Post-hoc confocal image showing the GCaMP6s (green) expression in TCs and MCs along with ChiEF-TdTomato^+^ GABAergic axons (red). Blue, DAPI. **b**, Pseudo-colored masks from MCs and TCs obtained by imaging in the MCL (top) and EPL (bottom). **c**, Example traces of MCs (red, top) and TCs (green, bottom) during light (blue shaded box), odors (orange contoured box), and odor and light simultaneous stimulation (blue shaded and orange contoured box). Scale bars, 5% ΔF/F and 2 s. **d**, Light-induced impact on MCs (red) and TCs (green) spontaneous activity (z-scored) in the presence (left) or absence (control, right) of ChIEF. Light illumination (2sec) was either pulsed (33 Hz) or continuous (CL). Ø: “shutter control”: closing/reopening the shutter without light stimulation. Violin plots are estimated ks density from the data. Black line, median; circle, individual cell. Light stimulation of ChIEF^+^ GABAergic cortical axons produced significantly greater inhibition than the respective control stimulation without ChIEF (ANOVA with Tukey’s post-hoc test, *** *P* << 0.001). In MCs, but not TCs, ChIEF CL stimulation produced greater inhibition than 33 Hz stimulation (two-sided t-test). Note that CL control in MCs is significantly greater than “shutter control” (two-sided t-test, *P*=0.02; *P*>0.05 for all other comparisons). **e**, Categorization (left) and magnitude (right) of the odor-evoked responses in MCs and TCs. Data are mean ± sem. **f**, MC (left) and TC (right) responses to simultaneous light and odor stimulation versus odor stimulation only (two-sided paired t-test). Circle, individual responsive cell-odor pair. White cross is mean ± s.d. for excitatory and inhibitory odor responses, separately. Solid line: equality line. **g**, In odor responsive neurons, light stimulation did not alter the intra-odor Euclidean distance (distance between the population responses to the same odor; “Within odor). However, In TCs, but not MCs, light stimulation increased the inter-odors Euclidean distance (distance between the population responses to the two odors, “Between odors”; two sided paired t-test).

Light stimulation of cortical GABAergic axons induced a significant reduction of spontaneous activity in the large majority of the MCs and TCs, both at 33 Hz and with CL (Fig. 6d; Supplemental Fig. 8a,b). In MCs, CL stimulation significantly reduced activity in a larger fraction of cells compared to 33 Hz stimulation, and inhibitory response magnitudes were larger. These differences were not observed in TCs. The observed inhibition was not an artifact of closing and reopening the PMT shutter. Indeed, in ‘shutter control’ trials, the number of cells exhibiting a significant change in activity was at statistical chance level, and the change in activity was significantly smaller, by an order of magnitude, than for light-stimulation trials (‘shutter control’ trials: trials with shutter closing, but no light presented, Fig. 6d; Supplemental Fig. 8b). We additionally controlled for an effect of blue light illumination *per se* in control animals that did not express ChIEF (‘light control’). A small, yet above chance proportion of MCs and TCs showed significant reduction of activity, consistent with previous reports (Ait Ouares et al., 2019), and that proportion increased with CL (Supplemental Fig. 8b). However, the magnitude of the light-induced inhibitory responses was 10-fold bigger in ChIEF-expressing animals compared to control animals not expressing ChIEF and therefore cannot significantly contribute to the reported effect (Fig. 6d).

GABAergic feedback inputs reduced spontaneous activity in the OB, but how does it influence incoming sensory feedforward information? Odor stimulation induced both inhibitory and excitatory responses in MCs and TCs, but with different relative proportions, as previously reported (Economo et al., 2016; Wachowiak et al., 2013; Yamada et al., 2017)(Fig. 6e). In odor-responsive cells, stimulation of GABAergic cortical axons induced a reduction of excitatory odor responses and a greater inhibition of inhibitory odor responses. This was true across both cell types and light stimulation patterns. The magnitude of the light-evoked inhibition and the odor responses were not correlated, resulting in linear subtraction of the odor-evoked activity (Fig. 6f). We also compared the impact of cortical GABAergic stimulation on spontaneous and odor-evoked activity at the individual neuron level. We found little correlation between the magnitude of the light-driven responses in spontaneous versus odor-evoked activity in both MCs and TCs (Supplemental Fig. 8c). For both cell types, light-driven inhibition was slightly stronger during spontaneous activity (Supplemental Fig. 8c).

To evaluate the effect of cortical GABAergic axon stimulation on the separation of odor representation within MC and TC populations, we calculated the Euclidean distance between population responses to either different (“Between odors”) or the same odor (Within odor”; Fig 6g). Consistent with a linear subtraction of the odor responses in MCs and TCs, light did not significantly alter the pairwise distance between population responses to a given odor (“Within odor” design; Fig 6g). In contrast, in the “between odors” design, light stimulation increased the distance in population odor representation of TCs, but not MCs (Fig 6g). This shows that stimulation of GABAergic axons specifically increases the difference in the representation of two different odors in TCs.

Another recipient of GABAergic projections in the OB is the internal part of the GL. We thus targeted our recordings to juxtaglomerular cells (JG cells, at the transition between the GL and EPL). As for MCs and TCs, CL stimulation of cortical GABAergic axons inhibited spontaneous activity of JG cells (Supplemental Fig. 8d). Odor stimulation drove mainly excitatory responses in JG cells, as reported previously (Banerjee et al., 2015) (Supplemental Fig. 8e). In the odor-responsive population, light stimulation of cortical GABAergic axons induced a linear reduction of the odor-evoked activity (Supplemental Fig. 8f). As seen in MCs and TCs, inhibition of spontaneous and odor-evoked activities was only weakly correlated (Supplemental Fig. 8f).

### Silencing cortical GABAergic outputs to the OB affects fine odor discrimination

Since GABAergic feedback modulates odor responses in OB output neurons, we next examined whether it could contribute to olfactory perception. To specifically inhibit cortical GABAergic feedback to the OB during the extent of an olfactory-guided task, we employed a pharmacogenetic approach. We expressed the inhibitory designer receptors exclusively activated by designer drugs (DREADD) hM4Di specifically in GABAergic AON/APC neurons. The axon terminals in the OB were selectively silenced by the local application of exogenous ligand clozapine-*N*-oxide (CNO), thereby sparing activity to other targets of the projections (Fig. 7a). The mice odor detection and discrimination thresholds were evaluated in a go/no-go task using carvone and limonene enantiomers (Fig. 7b). Detection threshold was assessed by diluting each day by a factor of 10 the two enantiomers to detect (from 1% to 0.0001% dilution). Discrimination threshold was assessed by presenting binary mixtures of the enantiomers, with a progressive and symmetric increase of the proportion of one into the other each day (from pure enantiomers discrimination, i.e., 100:0 vs. 0:100, to discrimination of mixtures with 55:45 vs. 45:55 enantiomer ratios).

**Figure 7.**
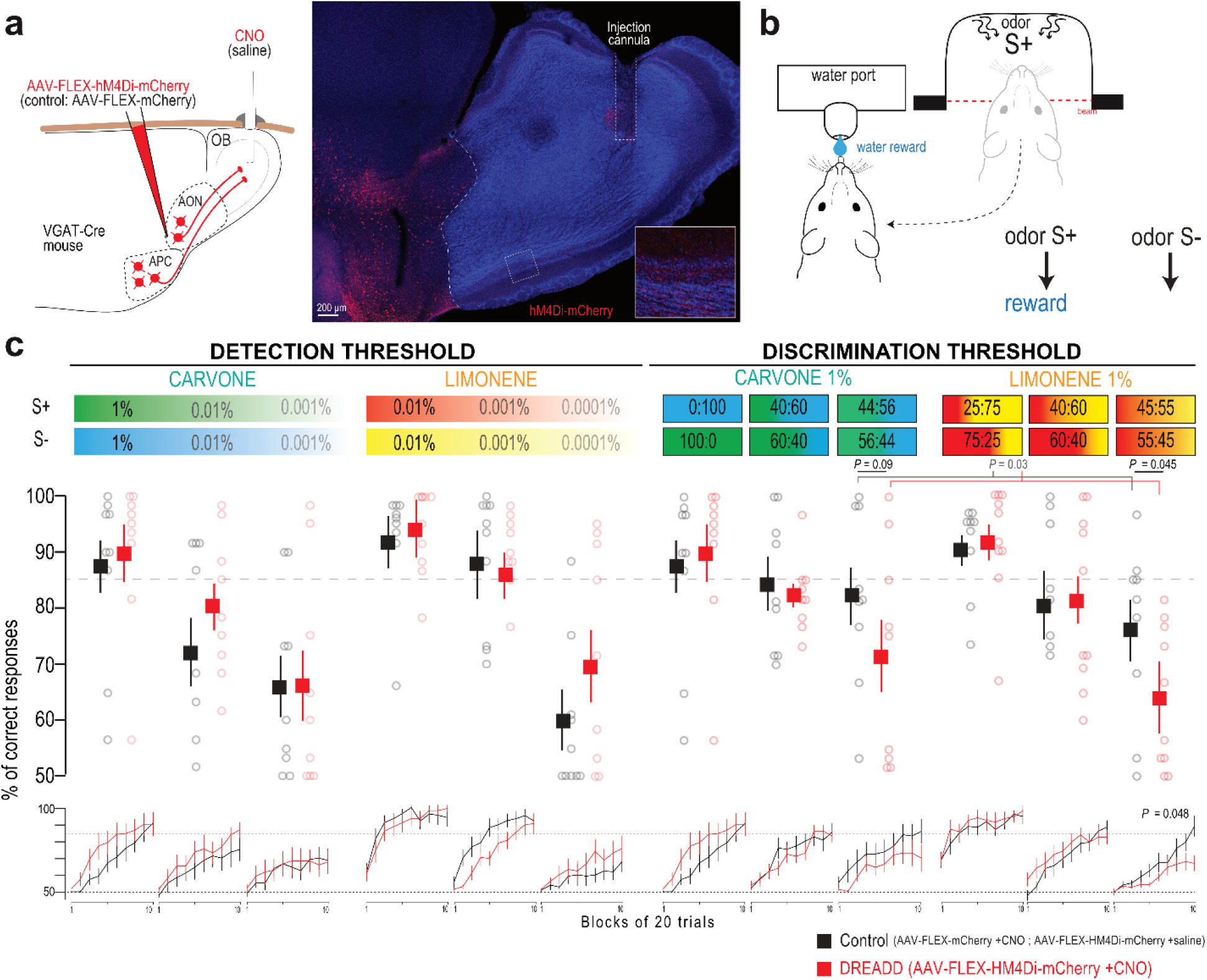
Targeted pharmacogenetic inhibition of cortico-bulbar GABAergic axons impairs fine odor discrimination. **a**, Specific silencing of axonal outputs of AOC GABAergic neuron expressing hM4Di-mCherry by locally injecting CNO in the OB (0.1 mg/mL, 1 µL per hemisphere). Control animal expressed the protein mCherry and received bilateral CNO injection. Right, Coronal section showing hM4Di-mCherry^+^ GABAergic cells in the AON and their axonal projections in the OB, together with the track of the injection cannula targeted to the core of the OB. Inset: high magnification of the boxed region with increased red fluorescence gain. **b**, Odor-reward association task. After a nose poke into the odor port, mice had to lick on the rewarded odor (S^+^) to obtain a water reward and refrained licking on the non-rewarded odor (S-). **c**, Performance (percentage of correct responses) for the discrimination of the carvone (green/blue) and limonene (red/yellow) enantiomers. Mean final performance (top; 3 last blocks, i.e., 60 last trials) and mean performance per block of 20 trials (bottom; 10 blocks per session, i.e., 200 trials). CNO reduced the performance for fine limonene mixture discrimination (enantiomers mixtures: 55/45 vs 45/55; top, two-sided Mann-Whitney test; bottom, repeated measure two-way ANOVA, F(9,153)=1.955, P=0.0483; hM4Di, n = 10 mice; Control, n = 9 mice). A similar trend was observed on carvone enantiomers, although it did not reach significance (n = 9 mice in each group, two-sided Mann-Whitney test). The effects of CNO on fine discrimination performance was still significant when analyzing together carvone and limonene performances (55/45 & 56:44, last three blocks; two-way ANOVA, F(1,33)=5.115, P=0.0304). Data presented as mean ± sem. Circle, individual mouse. Note that the data for carvone 1% and for carvone 100:0 are the same data.

Selective blocking of cortical GABAergic feedback had no effect on the detection of carvone or limonene enantiomers, even for very low odor concentration (Fig. 7c). In contrast, the discrimination of very similar binary mixtures of enantiomers was impaired. Indeed, a significant decrease in discrimination was observed for limonene enantiomers. A similar reduction in performance was observed for carvone enantiomers, although it did not reach statistical significance (Fig. 7c). When analyzing discrimination performances collectively for both pairs of enantiomers, blocking cortical GABAergic feedback did reduce fine odor discrimination performances (Fig. 7c). No significant difference in odor sampling time was observed (Supplemental Fig. 9). Altogether, the behavior data shows that silencing cortical GABAergic axon outputs to the OB impairs fine odor discrimination.

## Discussion

This study reveals the presence of GABAergic feedback projections from the primary olfactory cortex to the OB, with the AONp particularly densely packed with projecting neurons. We showed that cortical GABAergic feedback to the OB forms functional synapses with both GCL interneurons and principal output neurons – MCs and TCs. In awake mice, stimulation of this inhibitory feedback diminished both spontaneous and odor-evoked activities in GCL interneurons as well as in MCs and TCs. This global inhibition of the OB network was also captured by a computational model based on our experimental results. Interestingly, cortical GABAergic feedback separated odor population responses specifically in TCs, but not in MCs. Silencing of cortico-bulbar inhibitory axons altered performances during a fine odor discrimination task. Finally, we reported an analogous corticofugal inhibitory projection in the somatosensory system, suggesting a possible extension of our observations to other sensory systems.

As a first step to investigate the function of cortical GABAergic feedback in sensory systems, we manipulated these inputs collectively by labeling both the AON and APC, the two regions consisting of ∼95% of the cortical GABAergic OB-projecting neurons. Using genetics, immunolabeling and pharmacological tools, we showed that AOC GABAergic neurons contribute to the cortico-bulbar pathway. Using anterograde tracing, we show that GABAergic cortico-bulbar axons terminals 1) express the VGAT marker, 2) have a distinct laminar OB innervation profile compared to cortical glutamatergic or basal forebrain GABAergic projections, 3) do not result from a ‘leak’ in conditional viral expression or from a direct viral transduction of OB interneurons. We confirmed the existence of OB-projecting GABAergic neurons using four retrograde tracing methods (conditional HSVs, a double-conditional AAV approach, monosynaptic rabies tracing and conventional CTB-based retrograde). Electrophysiological recordings coupled with pharmacology confirmed that GABAergic AOC neurons form monosynaptic GABAergic synapse with OB neurons. The slightly longer IPSC latencies and slower kinetics in MCs and TCs (Fig. 3f, Supplemental Table 1) are consistent with input on electrotonically remote dendrites, presumably apical dendrites in the glomerular layer which are innervated by cortical GABAergic axons (Fig. 1b; Bardy et al., 2010). Lastly, data from our photometry and electrophysiological experiments — lack of GCL interneuron excitation and the different frequency recruitment of MC/TC population inhibition upon light-stimulation of cortical GABAergic versus glutamatergic axons — prove the specificity of our experimental approach and exclude a possible cross-reactivity with cortical glutamatergic axons.

OB-projecting GABAergic neurons originate from various olfactory cortical areas and express different neurochemical markers. We identified a dense cluster of GABAergic projection neurons in the AONp (at the border between the AONv, APC and OT). Early non-specific retrograde labeling studies had already identified a cluster of OB-projecting cells in the AONp in hamsters (Davis and Macrides, 1981; Davis et al., 1978), rats (Haberly and Price, 1978a; de Olmos et al., 1978) and mice (Diodato et al., 2016; Miyamichi et al., 2013; Shipley and Adamek, 1984; Zaborszky et al., 1986), yet their neurochemical content had not been specified. Recently, a study identified a cluster of lateral hypothalamus-projecting GABAergic neurons, presumably from the same region (coined ventral olfactory nucleus) (Murata et al., 2019), suggesting that the AONp could be a hub for broadcasting inhibition to olfactory and non-olfactory brain regions. Our data indicates that OB-and lateral hypothalamus-projecting GABAergic neurons are separate populations (Fig 2h) and further work should decipher the interplay between these two populations and whether they fulfill different functions.

Stimulation of cortical GABAergic feedback produced a net inhibition in both GCL interneuron and MC/TC populations. This finding is supported by a modeling approach where the reciprocally connected excitatory (MC/TC) and inhibitory subnetworks (GC) are both inhibited by GABAergic feedback stimulation — a phenomena akin to a “paradoxical” effect observed in cortical networks (Sadeh and Clopath, 2021). Our parsimonious network model also reproduces the frequency-dependent inhibition magnitude observed in our GCL and MC/TC fiber photometry data. We reasoned that upon weak cortical GABAergic feedback stimulation the reduction of the inhibitory drive from dSACs might counteract the direct inhibition from cortical feedback onto GCs. This interpretation is corroborated by work from Labarerra et al (Labarrera et al., 2013) showing that GCs are under tonic GABAergic inhibition in the awake state. Importantly, our model reproduces our experimental data with MC/TC-GC connectivity weight values producing physiological firing rates in both the excitatory and inhibitory subnetworks (low firing rate in GCs: (Cang and Isaacson, 2003; Cazakoff et al., 2014; Labarrera et al., 2013; Margrie and Schaefer, 2003); ∼15 Hz firing of MCs/TCs: (Fukunaga et al., 2012; Lepousez and Lledo, 2013; Mazo et al., 2016; Rinberg et al., 2006); Figure 5d).

At the functional level, we showed that silencing cortical GABAergic feedback axons disrupted fine sensory discrimination of similar odor mixtures, adding new evidence for a role of corticofugal projections in sensory detection and discrimination (Guo et al., 2017). Several non-exclusive mechanisms could account for the functional impact of cortical GABAergic feedback. First, GABAergic feedback facilitates beta band oscillations when stimulated at beta frequency. Beta oscillations have been shown to emerge during odor discrimination learning and require intact communication between the OB and the olfactory cortex (Lepousez and Lledo, 2013; Martin and Ravel, 2014). Moreover, precise spike timing of MCs and TCs relative to OB oscillations is critical for coding of odor intensity (Fukunaga et al., 2012; Shusterman et al., 2011; Smear et al., 2011, 2013), odor identity (Gschwend et al., 2012) and increases during olfactory learning (Li et al., 2015). Thus, altering the tightly regulated spike-field coherence could be a mechanism through which cortical GABAergic feedback directly shape odor discrimination. In the future, it would be interesting to address whether stimulating cortical GABAergic feedback at different phases of the sniff cycle differently impacts MC/TC activity and behavior. Second, cortical GABAergic feedback can modulate the time-window for integrating cortical excitatory inputs. Cortical glutamatergic axons drive excitation in MCs/TCs shortly followed by disynaptic inhibition (Boyd et al., 2012; Markopoulos et al., 2012), and GABA_B_ receptors activation specifically at cortical glutamatergic axon-to-GCs terminals can relax the temporal window for integration of excitatory inputs by MCs/TCs (Mazo et al., 2016). Cortical GABAergic axons could be the source of GABA activating GABA_B_ receptors. We reason that this heterosynaptic modulation of glutamatergic inputs could benefit from a fine temporal regulation in the cortex. Cortical GABAergic feedback is thus ideally positioned to modulate the timing for integrating cortical excitatory inputs (Grobman et al., 2018). Third, we reported that GABAergic feedback stimulation separated odor population responses specifically in TCs. Since MC and TC showed similar connectivity with cortical GABAergic inputs (Fig 3), differential impact on MCs and TCs could arise from differential intrinsic properties, local connectivity (Nagayama et al., 2014), or odor response properties (Burton and Urban, 2014; Fukunaga et al., 2012). Alternatively, MCs and TCs, or the GC populations they connect to (superficial vs. deep GC), could be connected distinctly to GABAergic cortical neurons (different cortical regions preferentially targeting MCs versus TCs and/or different types of GABAergic neurons). Either way, by separating TC population representation of different odors, GABAergic feedback stimulation can possibly enhance the discriminability capacity of a downstream decoder. Interestingly, in contrast to GABAergic feedback, manipulating glutamatergic feedback has been reported to alter the similarity of odor representation in MC, and not TC populations (Otazu et al., 2015). This suggests that GABAergic and glutamatergic cortico-bulbar projections may have distinct network effects and roles in olfactory behavior. Cortico-bulbar GABAergic projections also differ from basal forebrain GABAergic projections, the latter targeting specifically local GL and GCL interneurons (Hanson et al., 2020; Sanz Diez et al., 2019; Villar et al., 2021), resulting in a bidirectional modulation of MCs —switching from an inhibitory to disinhibitory net effect in the presence of odors— as well as a reduction of gamma oscillations (Böhm et al., 2020; Villar et al., 2021). Altogether, controlling the proper establishment of sensory-evoked network oscillations, modulating the time-window for cortically-driven excitation and separating representation of odor responses are three mechanisms through which GABAergic feedback can directly shape early sensory processing and odor perception.

Our observations highlight the advantage of direct cortical inhibitory projections. In cortico-thalamic and cortico-bulbar circuits, corticofugal glutamatergic projections produce disynaptic inhibition onto glutamatergic neurons through a GABAergic relay. In the paleocortex (olfactory system), this relay is mediated by local interneurons (Boyd et al., 2012; Markopoulos et al., 2012), while in the neocortex GABAergic relay neurons are located in the reticular thalamic nucleus (Crandall et al., 2015). In both paleo and neocortices, GABAergic relay neurons seem to implement band-pass filtering of the cortical glutamatergic drive. In the OB, stimulating cortical glutamatergic projections yields optimal inhibition of MCs and TCs in the beta range (20-40Hz) and decreases with faster stimulation regimes (Mazo et al., 2016). Similar frequency-dependent effect also takes place in thalamic nuclei: low-frequency cortico-thalamic axon stimulation suppresses thalamic activity while high-frequency stimulation enhances it (Crandall et al., 2015; Kirchgessner et al., 2020). In contrast, the strength of the inhibition driven by direct cortical GABAergic axons in MCs and TCs increased with increasing stimulation frequency and was faithful to even high frequency regimes (Supplemental Fig. 4d). Similarly, in the thalamus, direct extrathalamic GABAergic innervation from subcortical nuclei display a high fidelity to fast stimulation regimes (Halassa and Acsády, 2016). We also report cortico-thalamic GABAergic projection in the somatosensory system, suggesting that corticofugal GABAergic projections might be a common motif in sensory systems (Supplemental Fig. 4). Further studies will investigate whether this projection is also able to follow fast stimulation frequencies. In addition, our experimental data and OB network model showed that cortico-bulbar inhibitory inputs and their broad connectivity are in position to drive a global inhibition of the downstream network, limiting and preventing disinhibitory events on output neurons. Such a global network control triggers a synchronized reset of the network and may be engaged during specific brain states such as sleep (Halassa and Acsády, 2016).

## Material and Methods

### Animals

Adult (8-10 weeks at the time of injection) male and female VGAT-Cre (heterozygotes, Slc32a1^tm(cre)Lowl^, MGI ID: 5141270), SOM-Cre (Ssttm2.1(cre)Zjh, MGI ID: 4838416), VIP-Cre (Viptm1(cre)Zjh, MGI ID: 4431361), PV-Cre (Pvalbtm1(cre)Arbr, MGI ID: 3590684), Tbet-Cre (Tg(Tbx21-cre)1Dlc, MGI ID: J203355 (Haddad et al., 2013)) and C57BL/6JRj mice (Janvier Labs) were used in this study. This work was performed in compliance with the French application of the European Communities Council Directive of 22 September 2010 (2010/63/EEC) and approved by the local ethics committee (CETEA 89, project #01126.02, #2013-0086 and #DAP20025).

### Stereotaxic injections

Adeno-associated viruses (AAV) were generated by the Penn Vector Core, University of North Carolina Vector core, Addgene or produced by the Vector core of the Gene Therapy Laboratory of Nantes (INSERM UMR1089, https://umr1089.univ-nantes.fr/en/facilities-cores/cpv). Herpes simplex viruses (HSV) were produced by the MIT gene transfer core. CTB conjugated to Alexa Fluor 555 (C34776) was obtained from Molecular probes.

For viral injections, mice were deeply anesthetized using ketamine and xylazine mixture (150 mg/kg Imalgene and 5mg/kg Rompun, respectively; i.p.) and placed in a stereotaxic apparatus. A small craniotomy was performed, and a viral/CTB solution was injected into the brain through a glass micropipette attached to a Nanoinjector system (Nanoject II, Drummond). The coordinates and volumes used for injections were as follows: AON: 2.3 mm anterior and 1.1 mm lateral from Bregma, 3.3 and 3.6 mm deep from the brain surface, 100 nL/site; APC: 1.9 mm anterior and 2.25 lateral from Bregma, and 3.8 and 4.2 mm deep from the brain surface, 150-200 nL/site; NDB/MCPO: 0.1 mm anterior and 1.5 mm lateral from bregma, 5.5 deep from brain surface, 100nl/site; Somatosensory cortex S1, barrel field (BFC): 1 mm anterior and 3 mm lateral from Bregma, 1.2 and 1.5 mm deep from brain surface, 200nl/site. OB: 1mm anterior and 1mm lateral from junction of inferior cerebral vein and superior sagittal sinus, 1, 1.5 and 2mm deep from the brain surface, 100 nL/site. OB Lateral GL: 1 mm anterior and 2 mm lateral from junction of inferior cerebral vein and superior sagittal sinus, 1.5 mm from brain surface, 50 nL/site. Smaller volumes were used for sparse labeling and quantification of Synaptophysin-mRuby^+^ presynaptic boutons (Figure 1c): 50nL/site in the AON and 50 nL/site in the APC. Injection coordinates in the AON/APC were optimized to prevent any direct diffusion of the AAV virus in the OB (Supplemental Fig 1). Animals in which post-hoc histological examination showed that viral injections were not in the correct location were excluded from analysis. In the correctly injected animals, stereotaxic injection of AAV lead to virtually no somatas in the GCL (0.21 ± 0.11 neurons/mm^2^, n = 6 mice, 8 sections per mouse), as reported previously in wild-type mice (Lepousez et al., 2014).

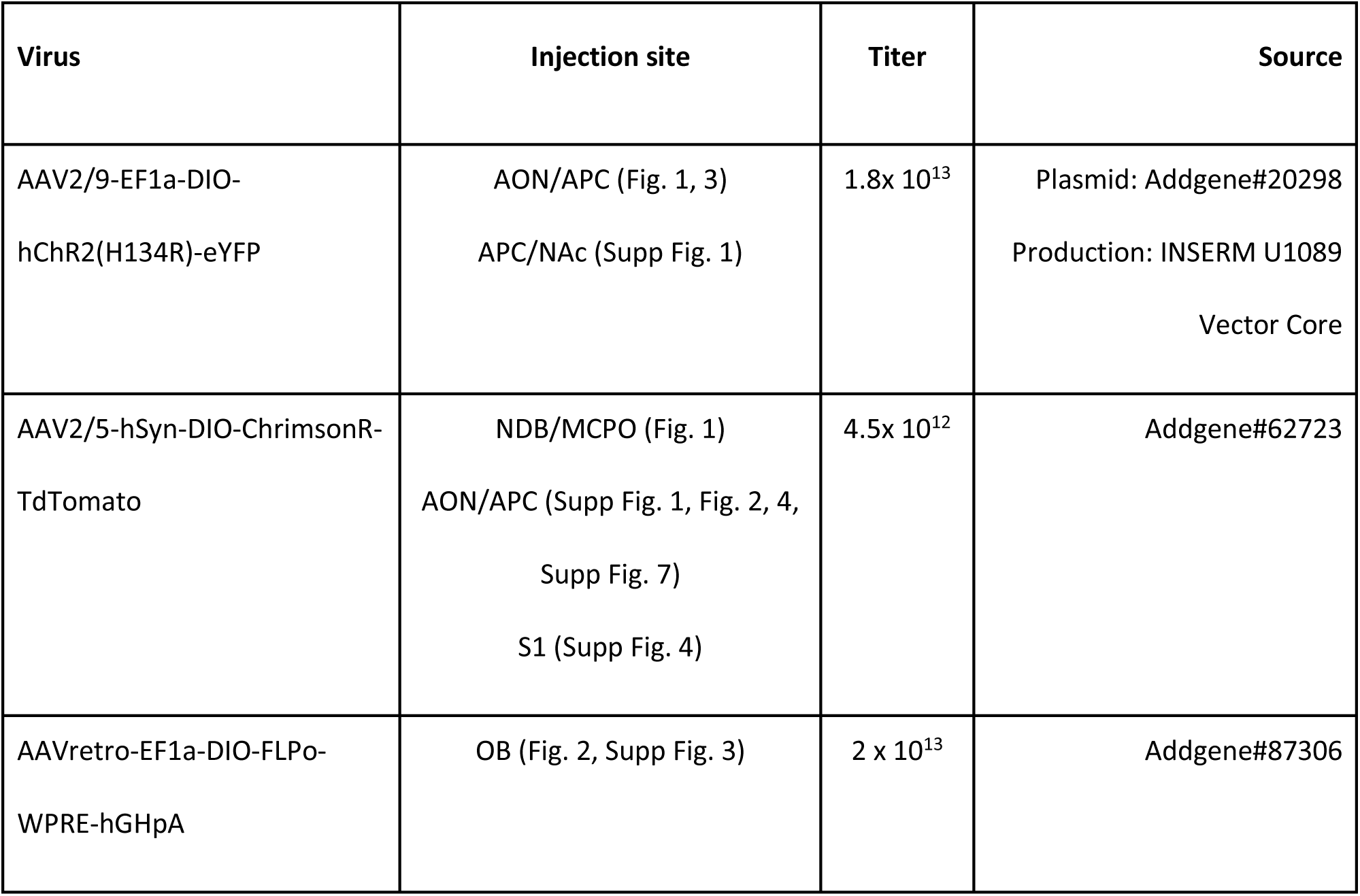
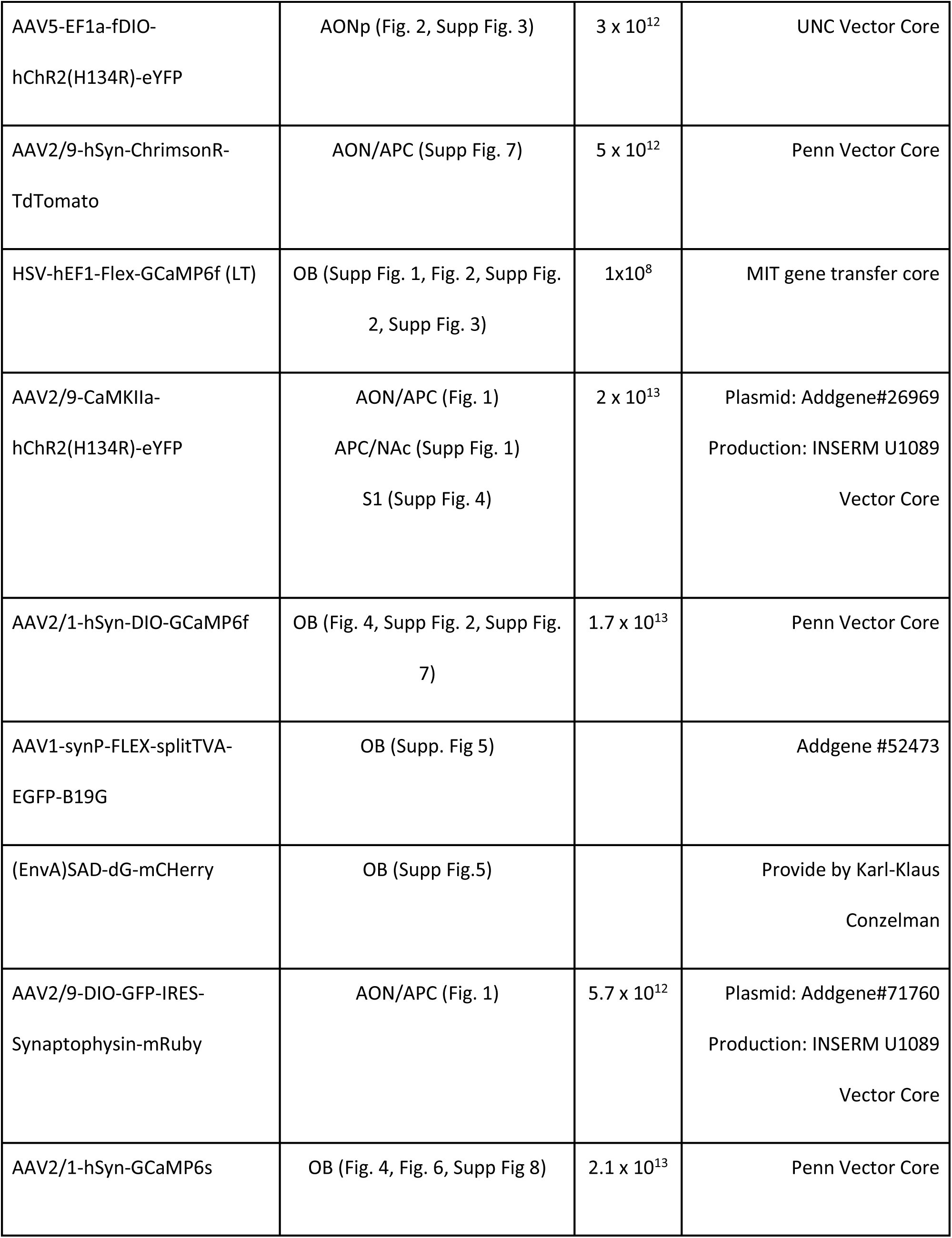
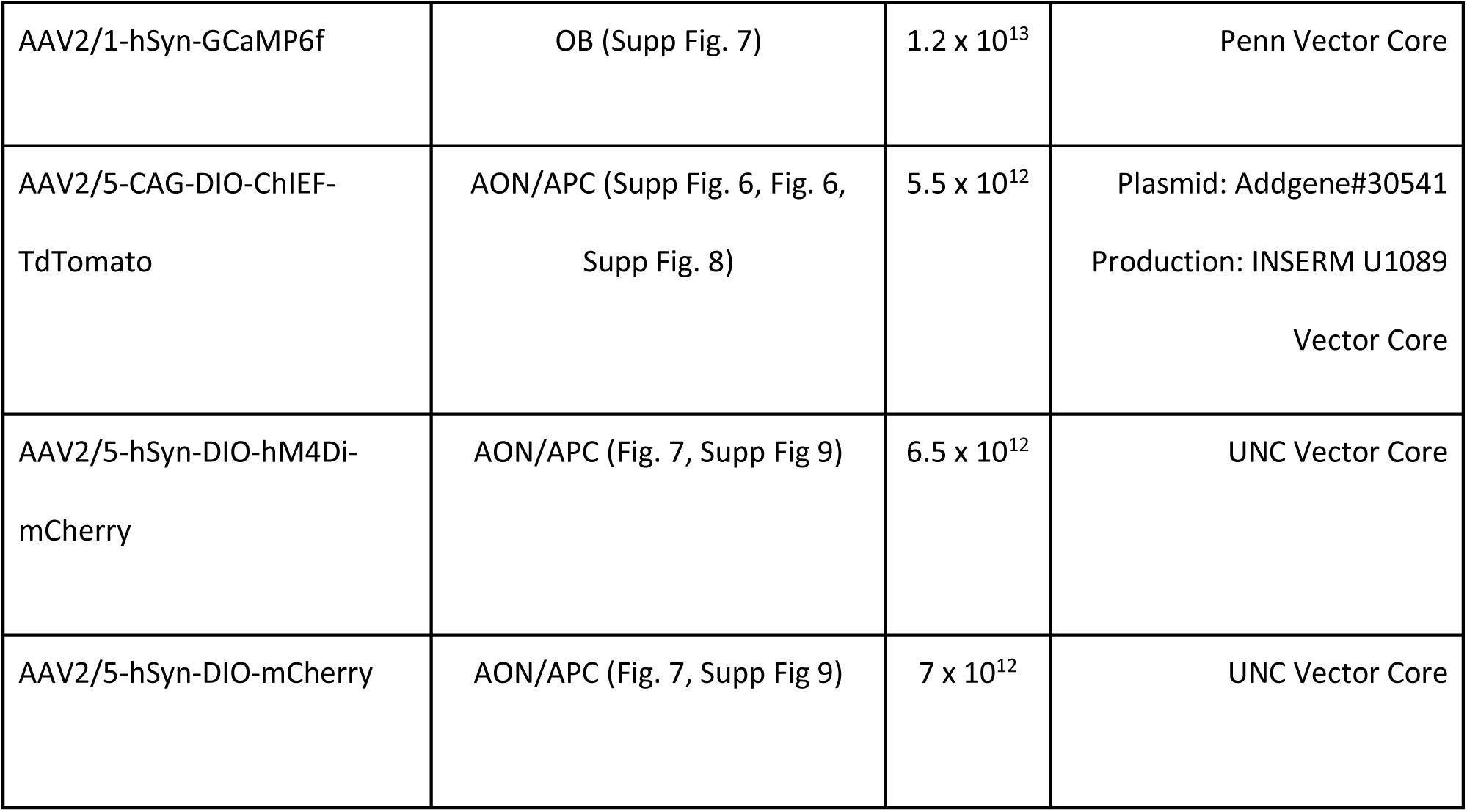

### Histology

#### Tissue preparation

Animals were intracardially perfused (4% paraformaldehyde (PFA) in 0.1M phosphate buffer) and the brains were removed and post-fixed in the same fixative overnight. Following cryoprotection (30% sucrose in PBS for 48h), brain sections were then cut with a freezing microtome (Leica). For post-hoc analyses of recording sites and viral expression, 60 µm-thick sections were sliced. OB sections were inspected to check for proper axonal expression, absence of virus diffusion into the OB, and for the absence of significant somatic labeling in the OB. 60 µm-thick sections were used for anatomical analyzes. For immunodetection of synaptic protein, animals were intracardially perfused (freshly prepared 4% paraformaldehyde (PFA) in 0.1M phosphate buffer), the brains were quickly removed, rinced in PBS and directly cryoprotected in (30% sucrose in PBS for 24h) without any post-fixation, before sectioning the OB on a freezing microtome (40 µm).

#### Immunohistochemistry

Primary and secondary antibodies used in this study are summarized in Table 2. Immunochemistry labeling was performed as follows: slices were rinsed, permeabilized and blocked in 10% Normal Goat Serum and PBS containing 0.25% Triton-X100 (PBST) for 2h. Primary antibodies were then incubated for up to 48h at 4°C in PBST containing 1% serum and 0.01% azide, washed three times and secondary antibodies were finally added for 2h in PBST containing 2% serum. Slices were then rinsed and counterstained with DAPI, mounted and imaged with a confocal microscope (LSM 700, Zeiss) or epifluorescence microscope (Axiovert 200, Zeiss) equipped with an Apotome system (Zeiss). For immunodetection of synaptic protein, the same protocol was used except the fact that PBST only contained 0.1% of Triton-X100 and primary antibodies were incubated for 72h. Confocal images (LSM 700, Zeiss) of the GCL were obtained with a 40X immersion objective (Zeiss) from the first 4µm from the slice surface, given the limited penetration of VGAT antibodies in such a GABAergic-rich region.

#### Cell counting of retrogradely-labeled cells

Coronal slices were serially collected and analyzed from the OB (+4 mm from Bregma) to the cortical amygdala (−0.5 mm from bregma). To evaluate cell density for each imaged slice, immunopositive somatas were manually counted for each subdivision and the surface of subdivision was measured (Zen, Zeiss). Counting was blind to the genotype in SOM-Cre, PV-Cre and VIP-Cre mice. Values for each subregion are averaged across sections for each mouse and used to calculate the mean cell density (averaged number of cells per mm^2^) and mean proportion of cells (% of cells counted in each subdivision relative to the total number of cells labeled in the respective brain). One out of every four slices were used for GFP counting in Figure 2c. One out of every six slices were used for GFP/SOM colocalization in Figure 2g.

#### Delineation of divisions and subdivisions of relevant brain regions

Brain regions were manually delineated using morphological parameters, DAPI staining, immunohistochemistry labeling, and the Allen Mouse Brain Reference Atlas (https://mouse.brain-map.org/static/atlas).

The *piriform cortex* is located in the ventrolateral forebrain, with a typical three-layered cortex and a layer 2 containing densely packed neurons. The piriform cortex was subdivided into anterior (APC) and posterior (PPC) regions, with the boundary at the caudal end of the lateral olfactory tract (LOT), as in (Mazo et al., 2017).

The *OT* is “readily identifiable as a large, pronounced, elliptical bulge nested between the LOT, the optic chiasm and the hemispheric midline ridge”. It is a “trilaminar region which contains a peculiar gyrating structure with anatomically defined ‘hills’ (gyri and sulci) and ‘islands’” (Wesson and Wilson, 2011). The OT stains heavily for choline acetyltransferase and this staining was used in some slices (Supplemental Figure 2b).

The *AON* is mainly located in the olfactory peduncle and consists of most of it. Yet, as detailed below, it extends caudally to the piriform cortex. The AON is a bilaminar region. Subdivisions of the AON were defined according to well-documented anatomical and cytoarchitectural landmarks (Brunjes et al., 2005, 2011; Haberly and Price, 1978b). The AON can be divided into two basic zone, the *pars externa* which is a “thin ring of cells that encircles the rostral end of the olfactory peduncle” (Brunjes et al., 2005) and the *pars principalis.* The latter is further subdivided in five regions, four of which are defined as a quadrant emerging from the anterior commissure: *pars dorsalis* (AONd), *pars ventralis* (AONv)*, pars lateralis* (AONl) *and pars medialis* (AONm). A fifth region extends caudally to the piriform cortex (*pars posterioralis*, AONp). AONl: area that lies directly under the LOT. AONl has the highest density of cells in the *pars principalis*, forming almost a visible layer. AONm, anterior section: AONm is lying below the OB. Posterior: AONm is delimited by the dorsal and ventral Tenia tecta. The ventral part of AONm also exhibits a cell-free gap which marks the border with AONv. AONd: facing orbito-frontal cortex, with no contact with LOT. The AONd is delimited on the medial border by the dTT. AONv: diffuse layer 2, with lower density of cells and no visible layer-like compared to AONl, which marks the border. AONv has limited contact with the LOT. The OT appears over the AONv in posterior sections.

AONp: caudal to AONv, buried between OT and APC, outside the olfactory peduncle per se. In contact with the anterior commissure, but with limited contact with LOT. The AONp starts when the dorsal and ventral TT fused and when the OT emerges clearly with 3 layers visible, isolating the AONp from the LOT. AONp has a group of large, loosely aggregated neurons (further references: (Davis et al., 1978; Macrides et al., 1981; Price, 1973).

The *NDB/MCPO* is a more caudal structure, located in the basal forebrain that runs rostro-caudally from the septum to the anterior amygdala area (de Olmos et al., 1978; Zaborszky et al., 1986).

#### Density of fluorescent axons

For the density profiles, OB coronal slices were imaged and the immunoreactivity profile was determined using ImageJ. Measurements were performed in matching slices and averaged across 4 sections per animal, normalized to the maximum intensity and then averaging between animals. For axonal density in Figure 2h, each section was imaged with the same parameter, and a binary image was calculated on ImageJ to then calculate the proportion of immunopositive pixels in each region. This value was then normalized to the density of immunopositive pixels in the injected region (AONp).

#### Quantification of synaptic punctas

The colocalization pattern of VGAT and mRuby immunoreactive clusters was determined in the GCL from 8-10 confocal sections per animal (n = 3 mice) using Imaris software (Bitplane, Zurich, Switzerland). For each section, each single-labeled VGAT^+^ or mRuby^+^ punctas was identified by Imaris surface segmentation algorithm (intensity threshold: 15% – 25% of the highest intensity value, surface threshold: 0.1 μm^2^) and the number and size of mRuby^+^ punctas was quantified. The colocalization algorithm was then used to identify the proportions of VGAT/mRuby colocalized punctas as well as the proportion of surface shared by the two markers. We considered a mRuby^+^ puncta co-labelled with VGAT^+^ punctas using a conservative estimate corresponding to a surface threshold of 0.1 μm^2^, i.e., 50% of the average mRuby surface.

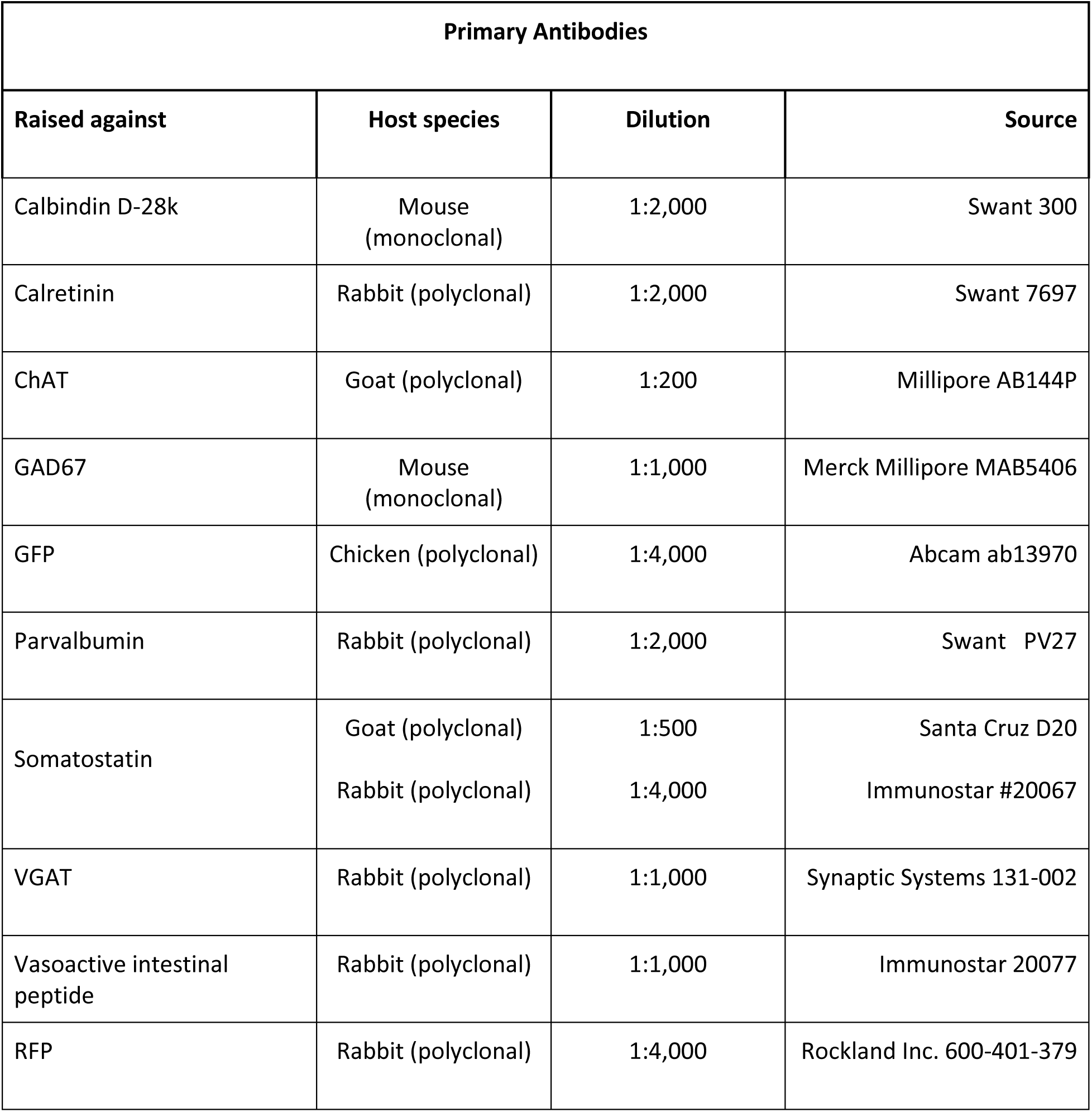
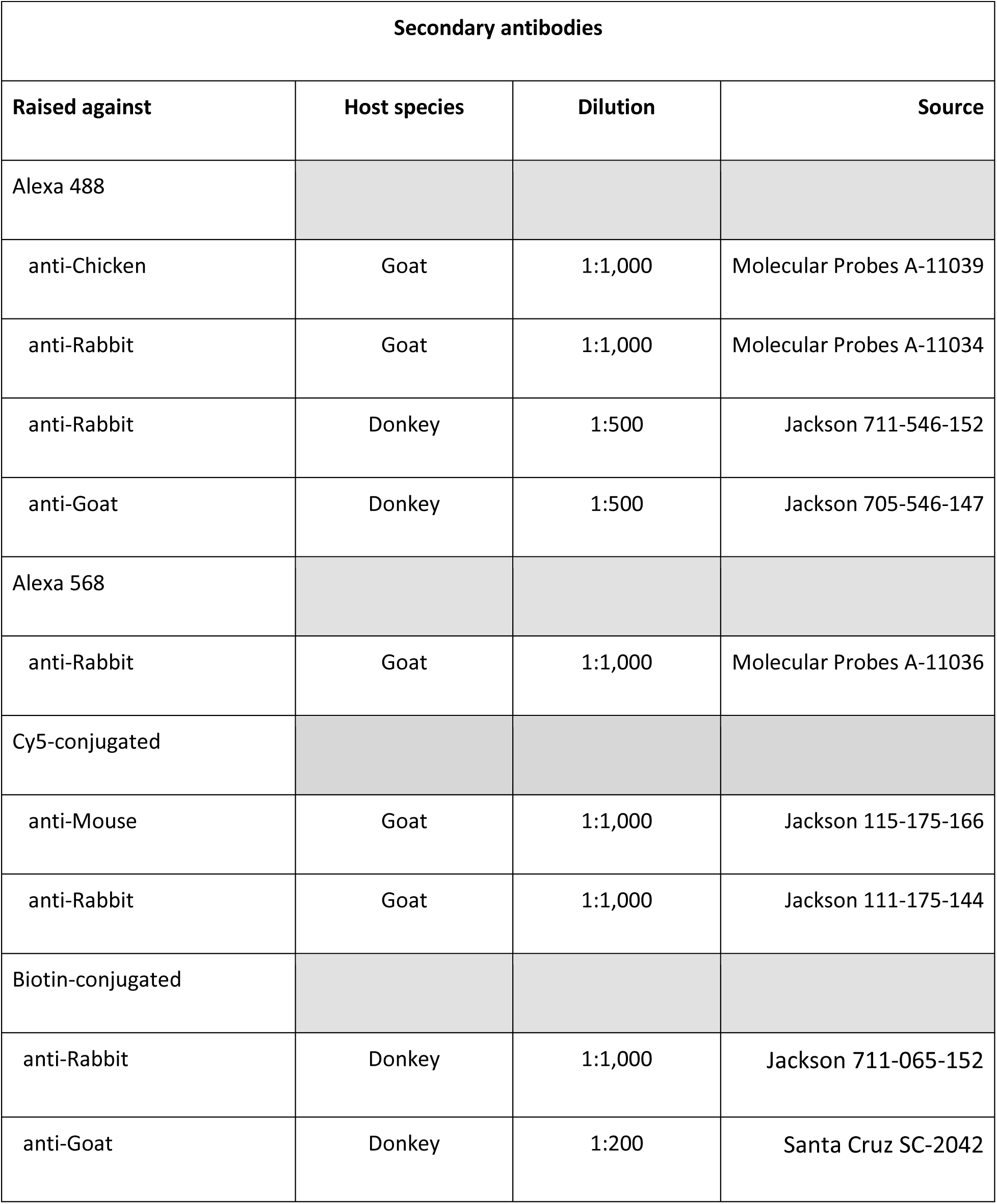

### Slice Electrophysiology

#### Slicing Procedure

Three weeks post injections, mice were deeply anesthetized with intraperitoneal injection of ketamine (100 mg/kg) and xylazine (10 mg/kg) and swiftly decapitated. The OB and frontal cortices were rapidly dissected and placed in ice-cold artificial cerebrospinal fluid (ACSF) containing 124 mM NaCl, 3 mM KCl, 1.3 mM MgSO4, 26 mM NaHCO3, 1.25 mM NaHPO4, 20 mM glucose, 2 mM CaCl [∼310 mOsm, pH 7.3 when bubbled with a mixture of 95% O2 and 5% (vol/vol) CO2; all chemicals from Sigma-Aldrich]. Horizontal slices (300-μm thick) of the OB were placed in bubbling ACSF in a warming bath at 35 °C for 30 min and then at room temperature (i.e., 22 ± 1° C). For whole-cell recordings, individual slices were placed in a chamber mounted on a Zeiss Axioskop upright microscope, and continuously perfused (1.5 mL/min) with 30°C ACSF (Warner Instrument inline heater). Slices were visualized using a 40× water immersion objective. Recordings were performed 3 weeks post-injection to avoid any possible contamination from adult-born GCs (Bardy et al., 2010).

#### Identification of neuronal subtypes

We obtained whole-cell patch-clamp recordings from visually targeted GCs, MCs, dSACs, PG cells, sSACs, eTCs and TCs. Neurons were filled with fluorescent dye (Alexa 488, 40 μM) and classified based on their somata laminar location, morphological, electrophysiological criteria (Boyd et al., 2012; Hayar et al., 2004; Murphy et al., 2005). Some patched cells were also filled with biocytin. For post-hoc revelation of biocytin-filled cells, slices were immediately fixed in PFA 4% for 24hrs, then rinced in PBS and incubated with PBS containing Triton (0.5%), DAPI (1:10000, Molecular Probes) and Alexa568-conjugated streptavidin (Molecular probes) for 2h. Slices were finally rinced, mounted (Fluoromount) and imaged with epifluorescence microscope (Axiovert 200, Zeiss) equipped with an Apotome system (Zeiss).

*eTCs*: large (∼20 μm diameter) somata in the GL, a single dendrite and tuft ramifying within one glomerulus, an axon extending into the EPL and a relatively low input resistance (∼200 MΩ).

*PG cells*: smaller somata (∼8-10 μm diameter) residing in the GL, with high input resistance (∼500-1000 MΩ). Highly ramified dendrite arbor in only one glomerulus.

*sSACs*: larger soma in the GL (> 10 μm diameter), low input resistance (∼200-300 MΩ), unique dendritic arbors that are exclusively periglomerular, span multiple glomeruli, lack tufts, and are poorly branched. *TCs*: large soma (10-20 µm diameter) in the inner part of the EPL (20-150µm above the MCL). Large apical dendrite innervating a single glomerulus, lateral dendrites in the EPL. Very low input resistance (∼50-100 MΩ)

*MCs*: very large soma in the MCL (20-30 µm diameter). Large apical dendrite innervating a single glomerulus. Lateral dendrites in the EPL. Very low input resistance (∼50-100 MΩ)

*dSACs*: soma size of > 10 µm diameter in the IPL or immediate surrounding GCL, with unique multipolar dendritic morphology (compared to GCs or MCs) and multiple neurites in the IPL. Low input resistance (∼200-300 MΩ).

*GCs*: small soma (8-10 µm diameter) in the GCL with one apical dendrite arborizing in the EPL and small basal dendrites, high input resistance (∼500-1000 MΩ). Patched GCs were preferentially located in the superficial GCL (100-150 µm below the IPL).

Averaged input resistances of patched cells are presented in Supplemental Table 1.

#### Recordings

Patch pipettes, pulled from borosilicate glass (OD: 1.5 mm, ID: 0.86 mm; Sutter instrument; P-87 Flaming/Brown micropipette puller, Sutter Instruments), had resistances of 6–10 MΩ for GCs and PG cells recordings and of 3-5 MΩ and were filled with a cesium gluconate-based solution: 126 mM Cs-gluconate, 6 mM CsCl, 2 mM NaCl, 10 mM Na-Hepes, 10 mM D-glucose, 0.2 mM Cs-EGTA, 0.3 mM GTP, 4 mM Mg-ATP, 280–290 mOsm, pH 7.3). Membrane potentials indicated in the text are corrected for a measured liquid junction potential of +10 mV. Recordings were obtained via an Axon Multiclamp 700B. Synaptic events were elicited by photo-activation of ChR2^+^ axon terminals stimulation using a 470-nm LED (Xcite by Lumen Dynamics) illuminating the sample through the objective. IPSCs were recorded at Vc = 0 mV, unless otherwise stated. Rise times were measured between 10% and 90% of peak amplitude. For IPSCs, decay time constants were derived by fitting the sum of two exponentials: F(t) = a × exp(−t/t fast) + b × exp(−t/t slow), where a and b are the peak amplitude of fast and slow components, respectively, and t fast and t slow are the respective decay time constants. Data were acquired using Elphy software (Gerard Sadoc, Centre National de la Recherche Scientifique; Gif-sur-Yvette, France) and analyzed with Elphy and IgorPro (Neuromatic by Jason Rothman, www.neuromatic.thinkrandom.com). In acute slices, none of the recorded GC exhibited ChR2-mediated inward currents (0/64) and IPSC kinetics in MCs and TCs were not consistent with GC-mediated inhibition (Bardy et al., 2010).

### In vivo electrophysiology

Following stereotaxic viral injection, a L-shaped metal bar and a silver reference electrode were fixed to the caudal part of the skull. Optic fibers [multimode, 430 µm diameter, numerical aperture (NA) 0.39, Thorlabs] were bilaterally implanted above the OB. Mice were allowed one week to recover and were subsequently slowly and progressively trained for head restraint habituation, a 5% sucrose solution was given as a reward. The craniotomy was performed the day before recording and protected with silicone sealant (KwikCast). An array of 4 tungsten electrodes (∼3 MΩ; FHC) was glued together and was slowly lowered into the OB. A drop of silicone sealant was applied to the brain surface to increase recording stability and avoid tissue desiccation. LFP signals were recorded in the MCL. Signals pre-amplified (HS-18; Neuralynx), amplified (1000 x; Lynx8, Neuralynx) and digitized at 20 kHz (Power 1401 A/D interface; CED). Light stimulation of cortical GABAergic axons was performed using an optic fiber coupled to a DPSS laser (473 nm, 150 mW; CNI Lasers; output fiber intensity, 20 mW) via a custom-built fiber launcher and controlled by a PS-H-LED laser driver connected to the CED interface. Light stimulation consisted in patterned light stimulation (10, 33 or 66 Hz) with 5-ms-long light pulses.

### Calcium imaging using fiber photometry

We used a fiber photometry system adapted from Gunaydin et al., 2014 (Gunaydin et al., 2014). Immediately following GCaMP6f virus injection in the OB, AON or APC, optical fibers (multimode, 430 µm in diameter, NA 0.5, LC zirconia ferrule) were bilaterally implanted close to the virus injection site, in the ventral part of the OB for GCL recording (1 mm anterior and 1 mm lateral from junction of inferior cerebral vein and superior sagittal sinus, 2 mm deep from the brain surface) and in the lateral part for MC/TC recordings (1 mm anterior and 1.5 mm lateral from junction of inferior cerebral vein and superior sagittal sinus, 1.5 mm deep from the brain surface) and then secured to the skull with a liquid bonding resin (Superbond, Sun Medical) and dental acrylic (Unifast). Three weeks post-injection, GCaMP6f was continuously excited using a 473 nm DPSS laser (output fiber intensity < 0.1 mW; Crystal Lasers) reflected on a dichroic mirror (452-490 nm/505-800 nm) and collimated into a 400 µm multimode optical fiber (NA, 0.48) with a convergent lens (f = 30 mm). The emitted fluorescence was collected in the same fiber and transmitted by the dichroic mirror, filtered (525 ± 19 nm) and focused on a NewFocus 2151-femtowatt photoreceptor (Newport; DC mode). Reflected blue light along the light path was also measured with another amplified photodetector (PDA36A, Thorlabs) for monitoring light excitation and fiber coupling. Red light (589 nm, 10 mW, pulse duration: 10-15ms) was collimated in the recording optic fiber to selectively activate cortical ChRimsonR-expressing GABAergic axon terminals in the OB while GCaMP6f was independently excited with low blue light intensity (< 0.1 mW), thereby avoiding cross-excitation of ChRimsonR (as in (Mazo et al., 2016)). Sessions with significant averaged changes in the reflected blue light (> 1% ΔF/F) were discarded from the analysis. Signals from both photodetectors were digitized by a digital-to-analog converter (DAC; Power 1401, CED) at 5 kHz and recorded using Spike2 software.

Mice were placed in a small, ventilated cage (∼0.5L). Using a custom-built air-dilution olfactometer controlled by the CED card, pure monomolecular odorants were diluted in mineral oil and saturated odorized air was further mixed with the air stream (1/10 dilution) before being delivered into the ventilated cage (flow rate of 4L/min), thanks to solenoid pinch valves. Odors were presented for 5 s every 60 s and dynamics of odor introduction and exhaust in the cage were constantly monitored using a mini photoionizer detector (miniPID, Aurora) positioned at the ceiling of the cage. Odors used were: Acetophenone 1%, Anisol 1%, Carvone+ 5%, Decanal 5%, Ethyl-butyrate 0.5%, Geraniol 5%, Heptanal 1% Hexanone 0.5%, 2-methyl-butyraldehyde 1%, Pentanol 1%, Valeraldehyde 0.2%, Methyl Salicylate 2%, 3-methyl-3-penten-2-one 1%. The 589 nm light stimulation was applied during 1 s, 3.5 s after odor onset when odor and light were simultaneously presented as well as 30s after odor presentation. For a given stimulation frequency, cycles of odor, light, and odor + light presentations were repeated 10 times for each condition to obtain interleaving trials with and without light (recording session duration ∼240 min). *S*ignals were smoothened (0.02 s window) and downsampled to 500 Hz. For each trial, the signal was normalized to the baseline fluorescence of the trial using the ΔF/F ratio with *F_0_* being the average fluorescence 2 s before the beginning of the trial. After completion of the recordings, mice were deeply anesthetized and transcardially perfused with 4% paraformaldehyde. OB and AOC were cut into 60μm-thick slices and observed with light and epifluorescence microscopes to evaluate the correct position of the optical fibers and the correct expression and diffusion of the virus. Animals in which post-hoc histological examination showed that viral injection or implanted optic fiber were not in the correct location were excluded from analysis. Selected sections were counterstained with DAPI and mounted for image acquisition (Axiovert 200 with Apotome system, Zeiss).

### Calcium imaging using two-photon microscopy

#### Acquisition parameters and imaging

After viral injections, a cranial window (3.0×1.4 mm glass) was placed over both OB and a stainless-steel head bar (L-shaped) was cemented to the skull. Mice were then allowed to recover for a month. During this period, the animals were progressively habituated to the head fixed position while staying quiet in the 50-ml open-ended support tube. Calcium activity was imaged using a two-photon system (950 nm, Spectra Physics) with a Prairie Investigator microscope (Bruker) and equipped with GaAsP photomultiplier tubes (PMTs). Ca^2+^ transients were imaged using a 16X, 1.05 NA microscope objective (Nikon) with a 2X digital zoom. The field of view was 512×512 pixels (423.7 x 423.7 μm), imaged at 15 Hz using a resonant galvanometer. Imaging planes (MCL, EPL or GL) were determined using anatomical landmarks and layers depth profiles as in (Adam et al., 2014; Sailor et al., 2016). Mean recording depth (relative to the GL) ± s.d: MCL, 201.8 ± 29.9μm; EPL, 60.1 ± 20.7μm. MCs were further distinguished from TCs by their denser packing and larger soma size (Otazu et al., 2015), less dense neuropil (Yamada et al., 2017).

#### Stimulation protocols

Trials consisted in 8s baseline, 2s stimulation (odor, light, or odor+light) and 10s inter-trial interval. Trials were grouped in blocks of 20 trials. 2-3 blocks were acquired per stimulus type. Data was acquired from 6 OBs of 4 animals.

##### Light activation of GABAergic cortical axons

The LED illumination for full-field photo-activation feature of the Investigator series (Bruker) was used to photo-stimulate the GABAergic axons in the OB. Blue light was directed to the field of view through the microscope objective. The PMT shutter remained closed during the photo-stimulation period and GCaMP6s fluorescence light was collected before and 50ms after the stimulation for allowing bidirectional realignment of the scanning. This time-window was evaluated using control trials where the light shutter closed but in the absence of photo-stimulation. GCaMP6s photo-bleaching using our ChIEF^+^ axon photo-activation paradigm was assessed by applying the same protocol to mice expressing GCaMP6s solely and was not minimal.

##### Odorant delivery

The odor pairs were a natural odor pair (curry powder vs. cinnamon) or a pair of pure monomolecular odorants (ethyl butyrate, valeraldehyde, isoamyl acetate, ethyl tiglate, hexanone or cineole, Sigma-Aldrich). Pure odorants were diluted 1:10 in 10mL mineral oil and natural odorants were presented in their native state. Saturated odor vapor was further diluted with humidified clean air (1:10) by means of computer-controlled solenoid pinch valves. Odor presentation was performed using a custom-built computer interface. Odor delivery dynamics were monitored and calibrated using a mini-PID (Aurora). Odors were delivered randomly within a block (10 trials of each odorant).

##### Odor and light stimulation

In “Odor + Light” trials, odorants and light were presented simultaneously utilizing the protocols mentioned above. After completion of the recordings, mice were transcardially perfused with 4% paraformaldehyde. OB and AOC were cut into 60μm-thick slices and observed with light and epifluorescence microscopes to evaluate the correct expression and diffusion of the virus. Selected sections were counterstained with DAPI and mounted for image acquisition (Axiovert 200 with Apotome system, Zeiss).

#### Image analysis

##### Motion correction

A full field of view motion correction was performed using a custom-made program in MATLAB. A two-dimensional cross-correlation of every frame with the average projection of the entire image set was used to identify the out of frame z-movements (Pearson’s r > 0.65 in the 2D cross-correlation). For lateral motion correction, the established ImageJ plugin *MoCo* (Dubbs et al., 2016) was used. In brief, it uses a Fourier-transform to improve the efficacy for identifying translational motion.

##### Principal component analysis (PCA) assisted reconstruction

We employed PCA on the raw motion-corrected datasets, after concatenating all the trials for each experiment into a 3-dimensional matrix. It leads to the possibility to express the original data in a lower dimension, capturing the largest variability in the dataset. The original image set is reconstructed from the most variable eigenvectors (non-varying (i.e., inactive) pixels and shot noise in the dataset do not have high variability across time and can be excluded). The reconstruction was done by a linear combination of the PC scores and the PC coefficients for the first 10 PCs of the dataset (Supplemental Fig. 10a,b).

##### Identification of regions-of-interest (ROIs)

The PCA-reconstructed images were used for the identification of ROIs. ROIs were manually drawn on the cell bodies using ImageJ and were imported in MATLAB (Supplemental Fig. 10c). As explained above, in order to remove the contribution of neuropil and background fluctuation, we performed a second PCA inside each ROI (Supplemental Fig. 10d-f). We used PC1-3 to redefine the outer bounds of each ROI for activity quantification. Note that PCA reconstructions did not modify the fluorescence data inside the ROIs. This process eliminated noisy signals (Supplemental Fig. 10g-m) and improved signal-to-noise ratio by 25% in a similar two-photon calcium imaging dataset (Saha et al., 2021).

#### Data analysis

*Z-score calculation*. For each ROI, the pixel intensities were smoothed across 5 frames and *z-score* was calculated for each cell as follow:

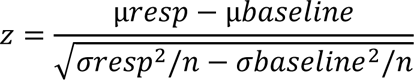

With µ and σ being the mean and standard deviation; resp, response (1 s after shutter reopening) and baseline is 1s before shutter closes. n is the number of trials. For comparison, we show in Extended Data Fig. 4 ΔF/F values, with F_0_ being the baseline determined for each trial.

Individual cell response to ChIEF light stimulation was considered significant if it passed a paired t-test based on single trials, with an alpha threshold of 0.01. Response and baseline values were the mean values 1s after and 1s before the shutter closed and reopened, respectively.

Odor responsive cells were identified using a two-sided paired t-test (α = 0.05) on all odor trials, regardless of whether GABAergic axons were stimulated or not to avoid a selection bias.

##### Euclidean distance

For each recording session (7 for MCs, 9 for TCs), pairs of population vectors were constructed from the averaged z-score responses to either the two odors presented on that day (Between odors design), or to the same odor (Within odor design). Only the cells responding to both odors (Between odors) or to the given odor (Within odor) were selected. Sessions were kept if a minimum number of 5 cells were responding to an odor. Pairwise Euclidean distance was calculated on the population vectors in the remaining sessions.

The Euclidean distance between two vectors *p* and *q* is given by the formula:

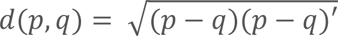

### Population Network Model

The firing rate of the excitatory (MCs or TCs, MC/TCs) and inhibitory subnetworks (GCs) were described by the following differential equations,

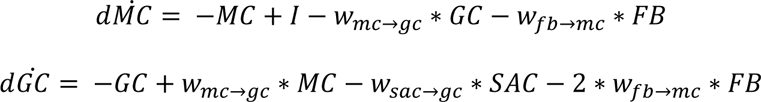

where *I* is the odor input, 𝑤_𝑚𝑐→𝑔𝑐_ is the strength of the connection between MCs (or TCs) and GCs and −𝑤_𝑚𝑐→𝑔𝑐_ is the reciprocal connection from GCs to MCs (or TCs). 𝑤_𝑠𝑎𝑐→𝑔𝑐_ is the strength of the inhibitory connection from dSACs to GCs. 𝑤_𝑓𝑏→𝑚𝑐_ is the strength of the cortical GABAergic feedback (FB) connection on MCs (or TCs). According to our slice electrophysiology results, the strength of the GABAergic feedback is roughly twice stronger on GCs, therefore it was set as 2 ∗ 𝑤_𝑓𝑏→𝑚𝑐_. The strength of the feedback on dSACs is even stronger and because, for the sake of simplicity, dSACs are not modeled, we modeled the inhibitory input strength to GCs in presence of feedback decreasing with the inverse of the strength of the feedback to MCs (or TCs).

The steady state (i.e., nullclines) of the MCs/TCs and GCs subnetworks are as following:

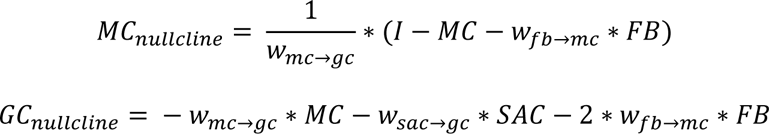

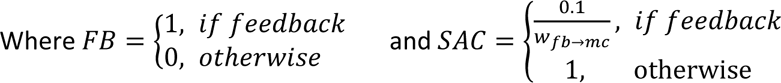

We explored different strengths and relative strengths between the MC-GC connections and the feedback.

The other parameters, such as the strength of the odor input and that of dSAC were not changed but they affect only the basal firing rate of MCs/TCs and GCs, not the slope of the nullclines nor the fixed points. Therefore, results will be qualitatively the same with different odor and dSACs input strength.

The fixed point (i.e., the equilibrium of the system) was used to determine the population rates of the MCs/TCs and GCs.

### Behavior

Two-guide cannulas (26-gauge, 7 mm long) were bilaterally implanted over the dorsal surface of the OB on the same day as viral injections 1mm anterior and 1mm lateral from junction of inferior cerebral vein and superior sagittal sinus. Guide cannulas were stabilized with a liquid bonding resin (Superbond, Sun Medical) and dental acrylic (Unifast) and a dummy cannula was positioned in the guide cannula to prevent blocking. Mice were habituated to be handled and maintained still while manipulating the dummy cannulas. On the day of the experiment dummies were retrieved, cannulas (8.5mm long, to inject at 1.5mm below the surface of the brain, 33-gauge and connected to a 10 μL Hamilton syringe) were placed for injections into the GCL. Dummies were put back in place a few minutes after the end of the injection. Behavior experiments were conducted using a go/no-go operant conditioning scheme as previously described. 2 weeks after the surgery, aged-matched adult male VGAT-Cre mice (10-12 weeks old) were partially water-deprived (maintained at 80-85% of their baseline body weight) and trained in custom-built computer-controlled eight-channel air-dilution olfactometers (Alonso et al., 2012; Lepousez and Lledo, 2013). Briefly, solenoid pinch valves controlled purified air streams, passing over the surface of mineral oil-diluted odorants. The odorized air was diluted 1:40 in odor-free air before its introduction into an odor sampling tube in the mouse operant chamber. Standard operant conditioning methods were used to train mice to insert their snouts into the odor sampling port for at least 1 s and to respond by licking the water delivery tube located 5 cm left of the odor port to get a water reward (3 µL). An infrared detection system continuously monitored the presence of the animal in the odor port. After this training phase to learn the procedure (200 trials per day for 5 days), mice had to learn to lick in the presence of a positive odor stimulus S+ and to refrain from licking and retract their head from the sampling port in the presence of a negative odor stimulus S-. In each trial, a single stimulus was presented and S+ and S- trials were presented in a modified pseudo-random order. Inter-trial intervals were minimum 8s-long. Each mouse performed a maximum of 10 blocks (200 trials) per day. The percentage of correct responses was determined for each block of 20 trials. A score of 85% at the very least implied that mice had correctly learned to assign reward/non-reward values. Odor sampling time was the time between the opening of the final valve and head retraction out of the odor sampling port.

Initial odor-reward learning, without intrabulbar injection, was performed using Anisole (S+) and Heptanone (S-). All the mice learned the behavioral procedure and were able to discriminate the two odors (behavioral performance > 85%) within three days. Three additional days of training were performed to ensure performance stabilization. Then mice were first trained with limonene enantiomers [S+, (+)-limonene; S-, (-)-limonene] and then carvone enantiomers [S+, carvone-(+); S-, carvone-(-)]. Detection threshold was assessed by diluting each day by a factor of 10 the two enantiomers to detect (from 1% to 0.0001% dilution). Discrimination threshold was assessed by utilizing binary mixtures of the enantiomers, with a progressive and symmetric increase of the proportion of one into the other each day (from pure enantiomers discrimination, i.e., 100:0 vs. 0:100, to discrimination of mixtures with 55:45 vs. 45:55 enantiomer ratios). To induce pharmacogenetic silencing before each different olfactory task, mice underwent bilateral intrabulbar injection of CNO or vehicle (saline) through the guide cannula (CNO final concentration: 0.1 mg/mL, injection speed: 0.33 µL/min for 3 min, 1 µL total/bulb) and were left in their home cage for 15-20 min to allow CNO or vehicle (saline) diffusion within the OB, before being placed in the olfactometer. The control group was composed of mice expressing mCherry in cortical GABAergic axons without the h4MDi receptor injected with CNO (controlling for CNO side-effects, n = 6) and mice expressing h4MDi in cortical GABAergic axons injected with saline (controlling for any non CNO-dependent side effect of expressing the exogenous h4MDi receptor, n = 3). For CNO injections, experimenters were blind relative to the viral constructs expressed in individual mice. For behavior, animals which did not perform the 200 trials in the 60 min time window following CNO injection were discarded from the analysis. After completion of the behavioral experiments, mice were transcardially perfused with 4% paraformaldehyde. OB and AOC were cut into 60μm-thick slices and observed with light and epifluorescence microscopes to evaluate the correct position of the injection cannula and the correct expression/diffusion of the virus. Animals in which post-hoc histological examination showed that transgene expression were not restricted to the AON were excluded from analysis. Selected sections were counterstained with DAPI and mounted for image acquisition (Axiovert 200 with Apotome system, Zeiss).

### Statistical analysis

Sample sizes are indicated in the figure and/or in the legend of the corresponding figures. All statistics were performed using GraphPad Prism 8 or MATLAB. Data containing two experimental groups were analyzed using unpaired two-sided Student’s t-test (parametric observations), unpaired two-sided Mann– Whitney test (non-parametric observations), one-way and two-way ANOVA tests followed by Tukey’s post hoc analyses to account for multiple comparisons. Data containing multiple paired measures were analyzed using repeated-measures ANOVA test. The mean and s.e.m. are reported for each experimental group.

## Acknowledgements

We thank Dr. William Podlaski for help with the model. We thank Sara Moberg for comments on the manuscript. We also thank Carine Moigneu for viral injections, Julien Grimaud and Lucie Dixsaut for early works. We also wish to thank Uwe Maskos and David DiGregorio from the Institut Pasteur for the gift of SST-Cre, PV-Cre and VIP-Cre mice. We also thank the Genetically-Encoded Neuronal Indicator and Effector (GENIE) Project and the Janelia Farm Research Campus of the Howard Hughes Medical Institute for sharing GCaMP6f constructs. We are greatful to Alexandru A. Hennrich and Karl-Klaus Conzelmann (Max Von Pettenkofer Institute Virology and Gene Center, Medical Faculty, Ludwig-Maximilians-University Munich, Germany) for the generous gift of (EnvA)SAD-ΔG-mCherry virus. We also thank the Vector Core of the Laboratory for Translational Research in Gene Therapy (INSERM UMR 1089, Université de Nantes) for AAV vector production and the Viral Core Facility of the McGovern Institute (MIT) for HSV production. This work was supported by the life insurance company “AG2R-La-Mondiale”, the Agence Nationale de la Recherche (ANR-15-CE37-0004 “SmellBrain’’ and ANR-16-CE37-0010 “ORUPS”) and the Laboratoire d’Excellence Revive (Investissement d’Avenir, ANR-10-LABX-73). Our laboratory is part of the Ecole des Neurosciences de Paris (ENP) Ile-de-France network and is affiliated with the *Bio-Psy* Laboratory of Excellence. C.M. is a recipient of a fellowship from the French Ministère de l’Education Supérieure et de la Recherche and was also supported by the Fondation de la Recherche Médicale (FDT20160435483).

## Author contributions

C.M., G.L., A.N., S.S. and P.-M.L. designed the experiments, C.M., G.L., A.N., S.S., and E.P. performed and analyzed the experiments. C.M., G.L., S.S. and P.-M.L. wrote the manuscript.

## Data availability

The datasets generated during the current study as well as the custom MATLAB code used to analyze the data are available from the corresponding authors upon reasonable request.

---

**Supplemental Fig. 1.**
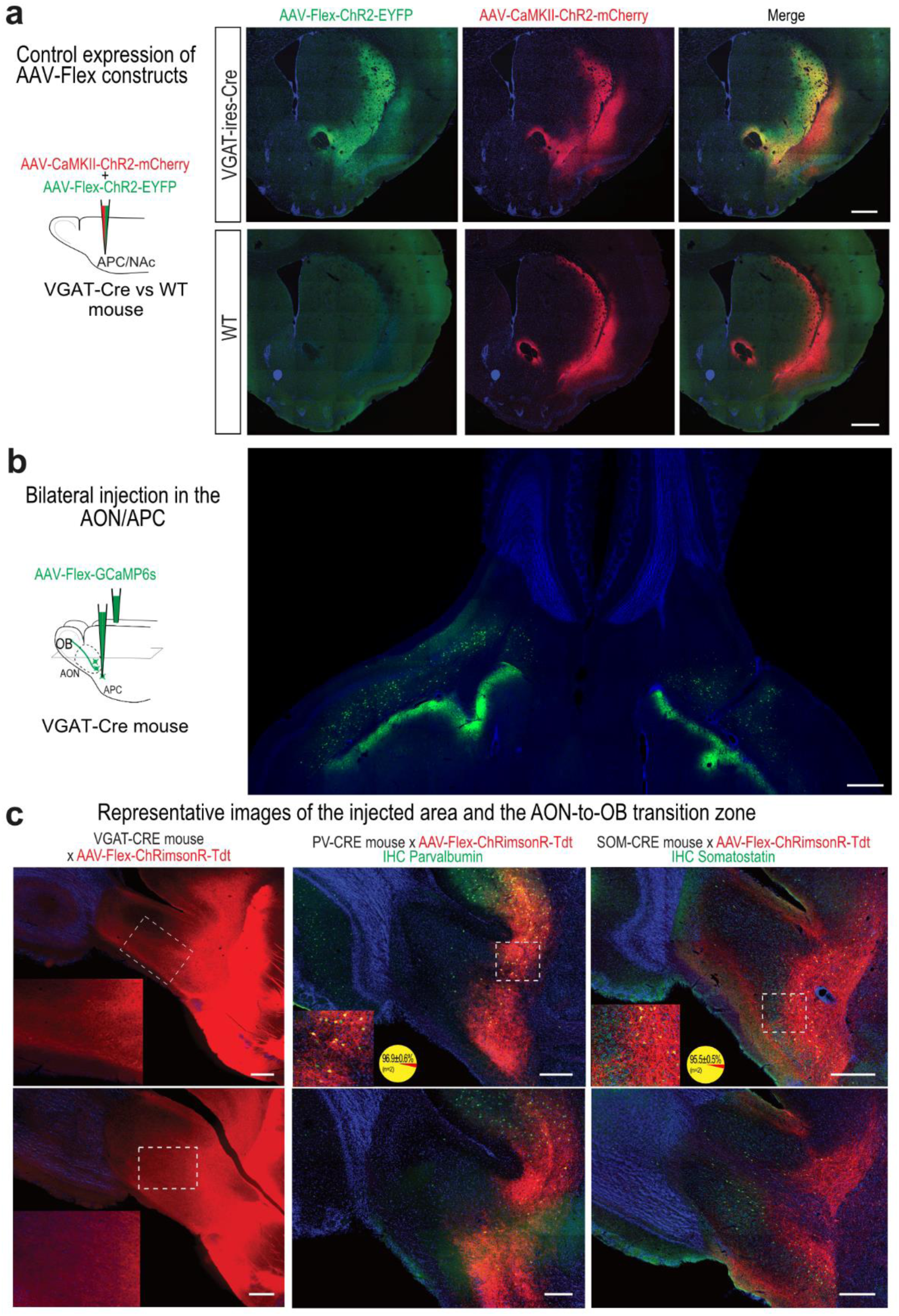
Further characterization of the anterograde labeling. **a**, Cre-dependent virus specificity control. Injection of a mix of Cre and non-Cre dependent viruses (AAV-Flex-ChR2-eYFP and AAV-CaMKII-Chr2-mCherry, respectively) were injected in VGAT-Cre versus WT mice. Three weeks post injection, no labeling associated with the Cre-dependent virus was observed in WT mice. Co-injection of the non-Cre dependent virus assured the proper targeting of the injection. Injections were targeted in the APC as well as in the nucleus accumbens (NAc) as the latter contains a high proportion of GABAergic neurons compared to the cortex. Lack of neuronal expression in the nucleus accumbens supports the tropism of the CaMKII promoter for excitatory neurons. **b,** Horizontal slice showing the bilateral injection sites in the AON/APC of VGAT-Cre mice. **c,** Representative images of the injection site in VGAT-Cre (left), PV-IRES-Cre (middle) and SOM-IRES-Cre (right) mice. Top and bottom rows are different parasagittal planes of the same mice. Bottom left are insets of the dashed boxes in the same image. In VGAT-Cre mice, the virus only invaded the posterior part of the peduncle and did not diffuse in the OB. Note the lack of neurons infected in the SVZ (top). We confirmed the specificity of the mouse lines and the efficiency of our IHC staining protocols against PV and SOM markers in PV-IRES-Cre and SOM-IRES-Cre mice, respectively. Colocalizations percentages are percentages of virally-labeled cells that were co-labeled with IHC for the corresponding marker (n = 2 mice). Scale bars are 500 µm (**a, b**) and 250 µm (**c**).

**Supplemental Fig. 2.**
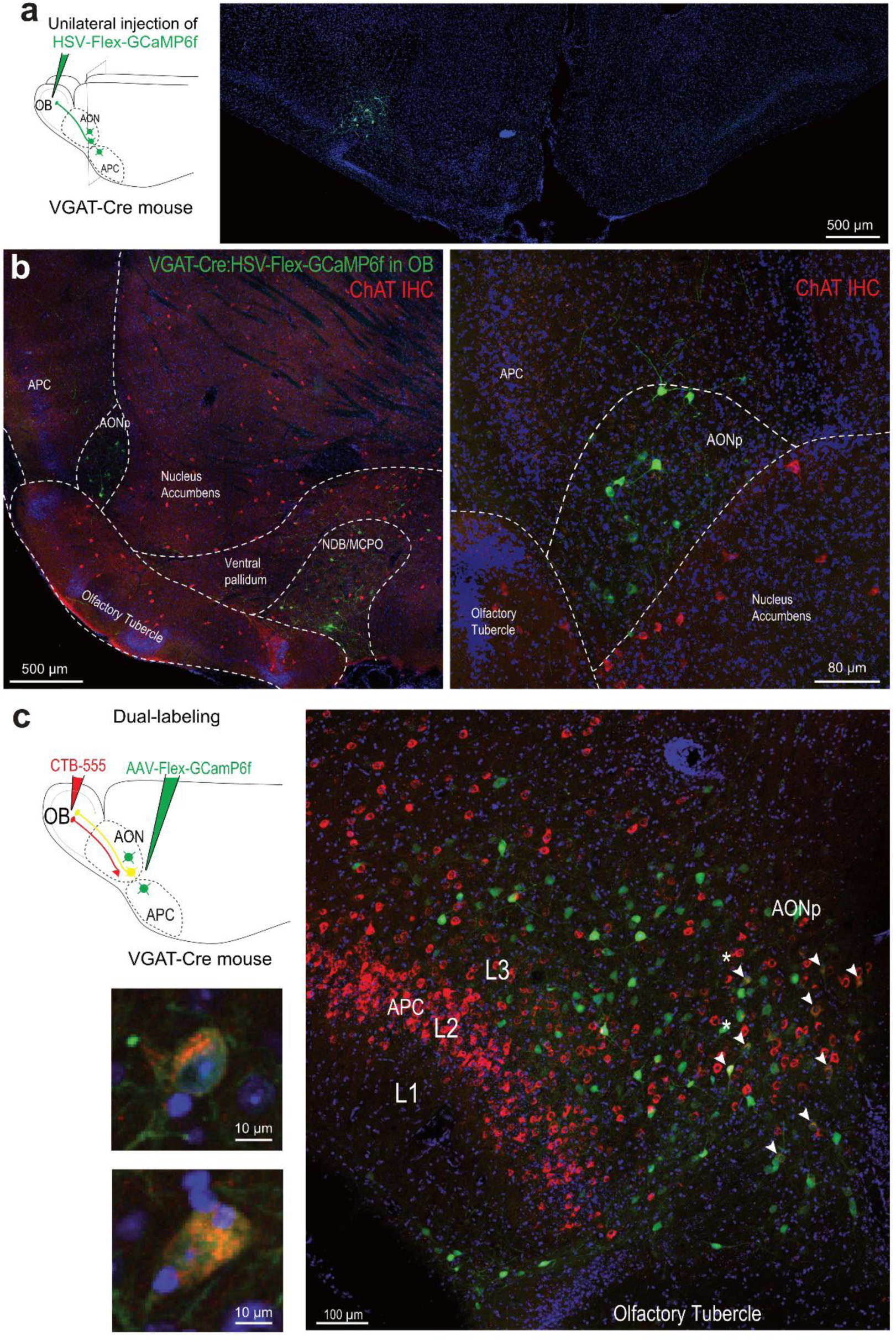
Further anatomical and neurochemical analysis of the OB-projecting GABAergic cells in the AON/APC. **a,** Left, Unilateral OB injection of the retrograde virus HSV-Flex-GCaMP6f labeled OB-projecting GABAergic neurons ipsilaterally, but not contralaterally in the AON and APC. Right, coronal slice through the AONp. **b**, Choline acetyltransferase (ChAT) IHC in a AONp-containing brain section showing that OB retrogradely-labeled GABAergic cells are located outside the cholinergic-rich brain area (compare with cells in the NDB/MCPO). This argues against the hypothesis that this region is a rostral extension of a striatal or pallidal structure. **c**, Non-selective retrograde labeling of OB-projecting cells (CTB-555) and somatic viral labeling of GABAergic neurons recapitulate the observations with HSV. Bottom, coronal slice through the APC/AONp (1.7 mm anterior to Bregma). CTB produced a classic labeling of cortico-bulbar glutamatergic cells in L2 and L3 in the APC. Double-labeled cells (arrowheads) were mainly found in the AONp, located at the border between the APC and OT. Starred cells are magnified in the top panels. Blue, DAPI.

**Supplemental Fig. 3.**
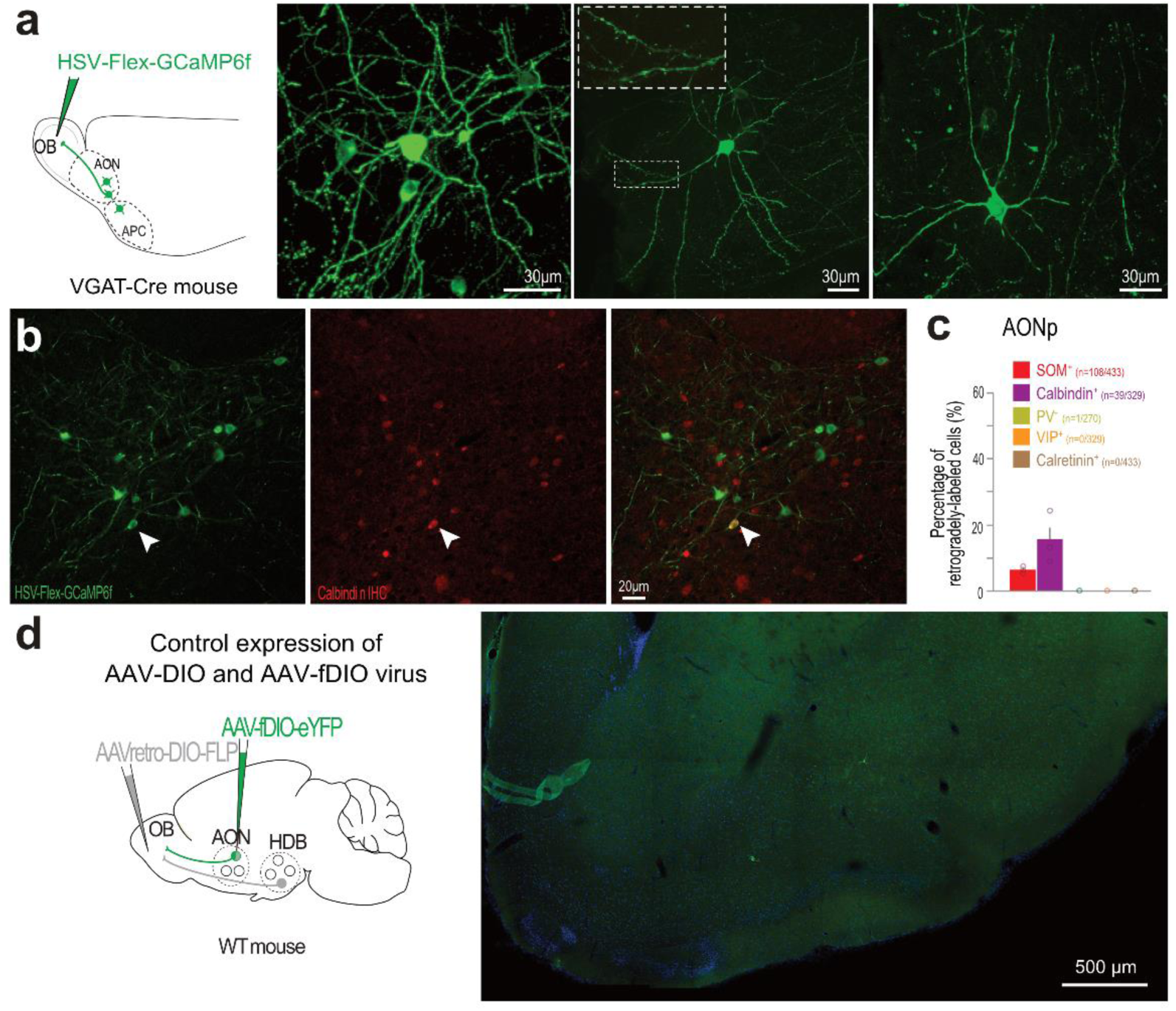
**a**, Magnification of examples of retrogradely-labeled cells. Middle and right panels show spiny (see inset, detail of the boxed region) and unspiny neurons from the AON/APC. **b,** IHC for calbindin labeled OB-projecting GABAergic neurons (VGAT-Cre x HSV-Flex-GCaMP6f in the OB) in the AONp (arrowhead). **c,** Percentage of the OB-projecting GABAergic neurons in the AONp that expressed classical markers for GABAergic neurons (n = 3 mice). Data is mean ± sem across mice. Circles are individual mice. n numbers are the number of double-labeled cells counted for each molecular marker amongst the total analyzed retrogradely-labeled cells. **d,** Specificity control for our INTRSECT labeling. Same as figure 2h, but in wild-type mice. A single neuron was labeled in the entire AOC (right).

**Supplemental Fig. 4.**
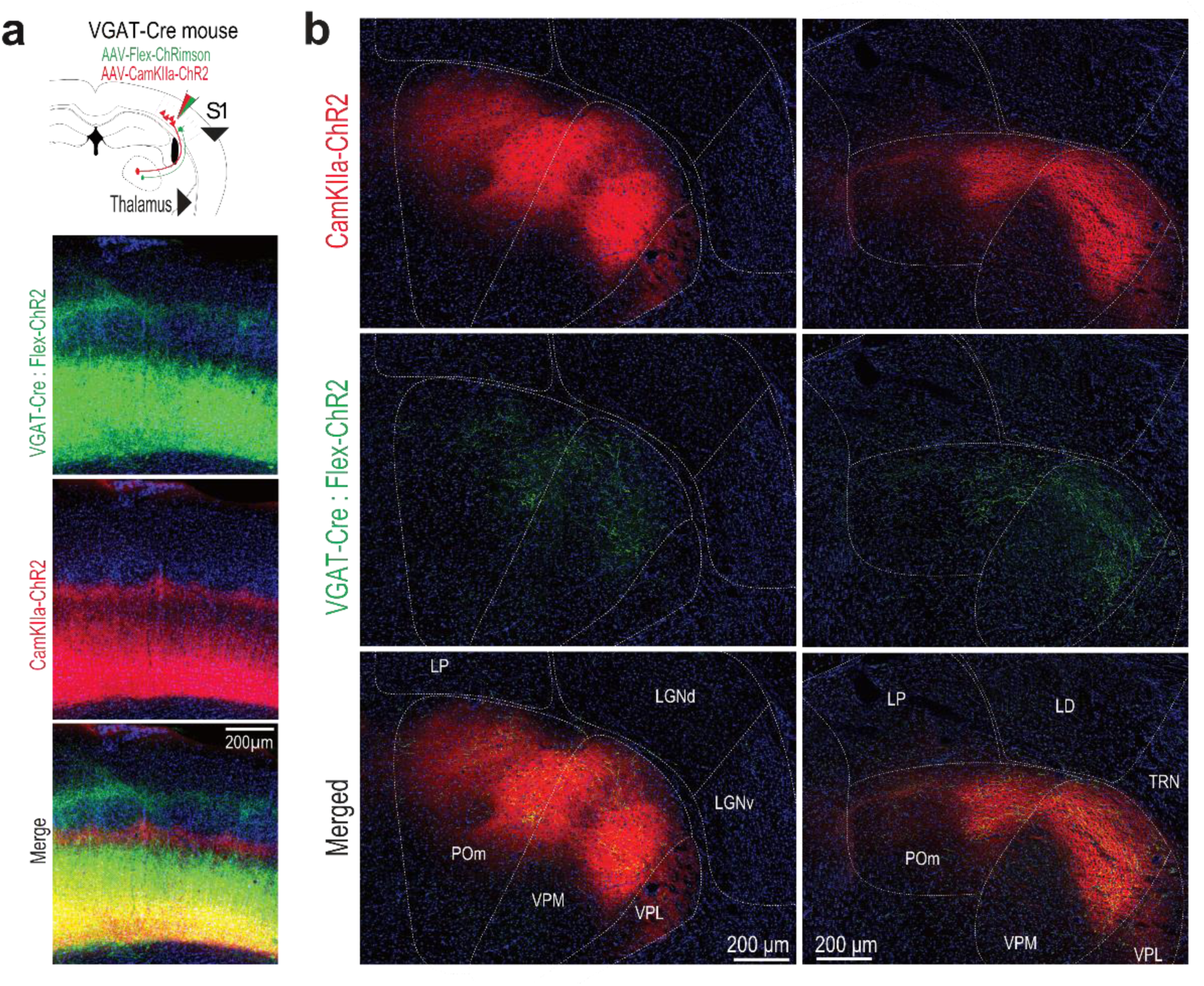
The primary somatosensory cortex (S1) also sends GABAergic projections back to the somatosensory thalamus nuclei. **a**, Anterograde labeling of S1 barrel field glutamatergic (ChR2-mCherry) and GABAergic (ChR2-eYFP) axons in the sensory thalamic nuclei. Right, Injection site in L5/6 of S1, barrel field. Blue, DAPI. **b**, Glutamatergic (red) and GABAergic (green) axons across 2 sections from 2 mice through the sensory thalamic nuclei. Blue, DAPI. Thalamic nuclei: LD: latero-dorsal LP: lateral posterior; POm: posteriomedial; VPL: ventral posterolateral; VPM: medial ventral posteriomedial; TRN: thalamic reticular nucleus.

**Supplemental Fig. 5.**
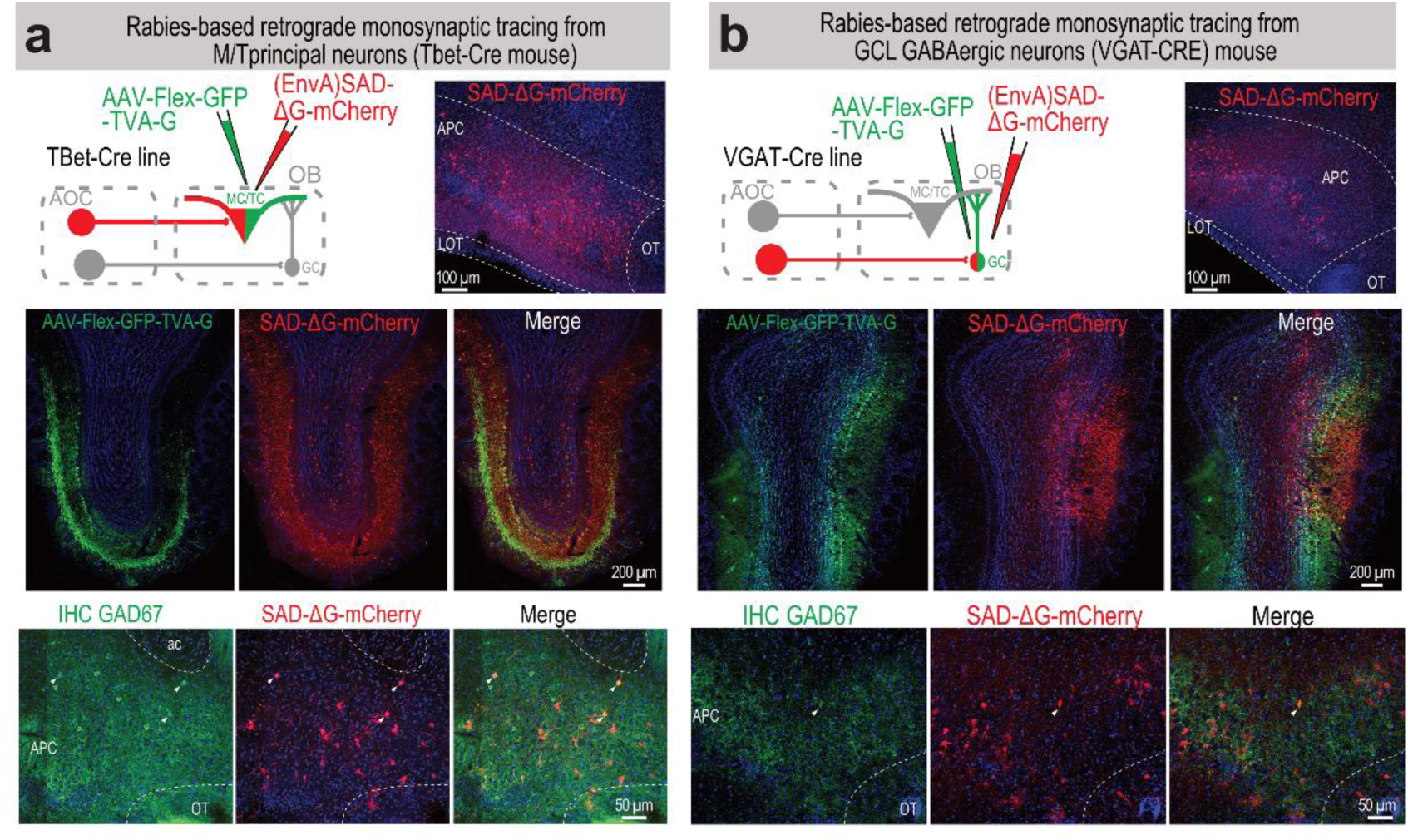
Monosynaptic retrogradely-labeled neurons from genetically identified OB neuron populations. **a**, Rabies-based transsynaptic retrograde tracing from MCs and TCs (“starter cells”). An AAV encoding the Cre-dependent GFP, avian virus receptor (avian tumor virus receptor A, TVA) and rabies glycoprotein (G) was injected into the OB of Tbet-Cre mice that express Cre recombinase specifically in MCs and TCs. Subsequent injection of G-deleted envelope protein from avian ASLV type A (EnvA)-pseudotyped rabies virus ((EnvA)SAD-ΔG-mCherry) into the OB resulted in transsynaptically retrogradely labeled cells in the olfactory cortex (top, right). IHC for GAD67 revealed that some AOC neurons projecting onto MCs and TCs are GABAergic (bottom, arrowheads). **b,** Same as **a** but with GCL interneurons as starter cells (injections in the GCL of VGAT-Cre mice). Some double-positive cells were sparsely found in the AOC (bottom, arrowhead).

**Supplemental Fig 6.**
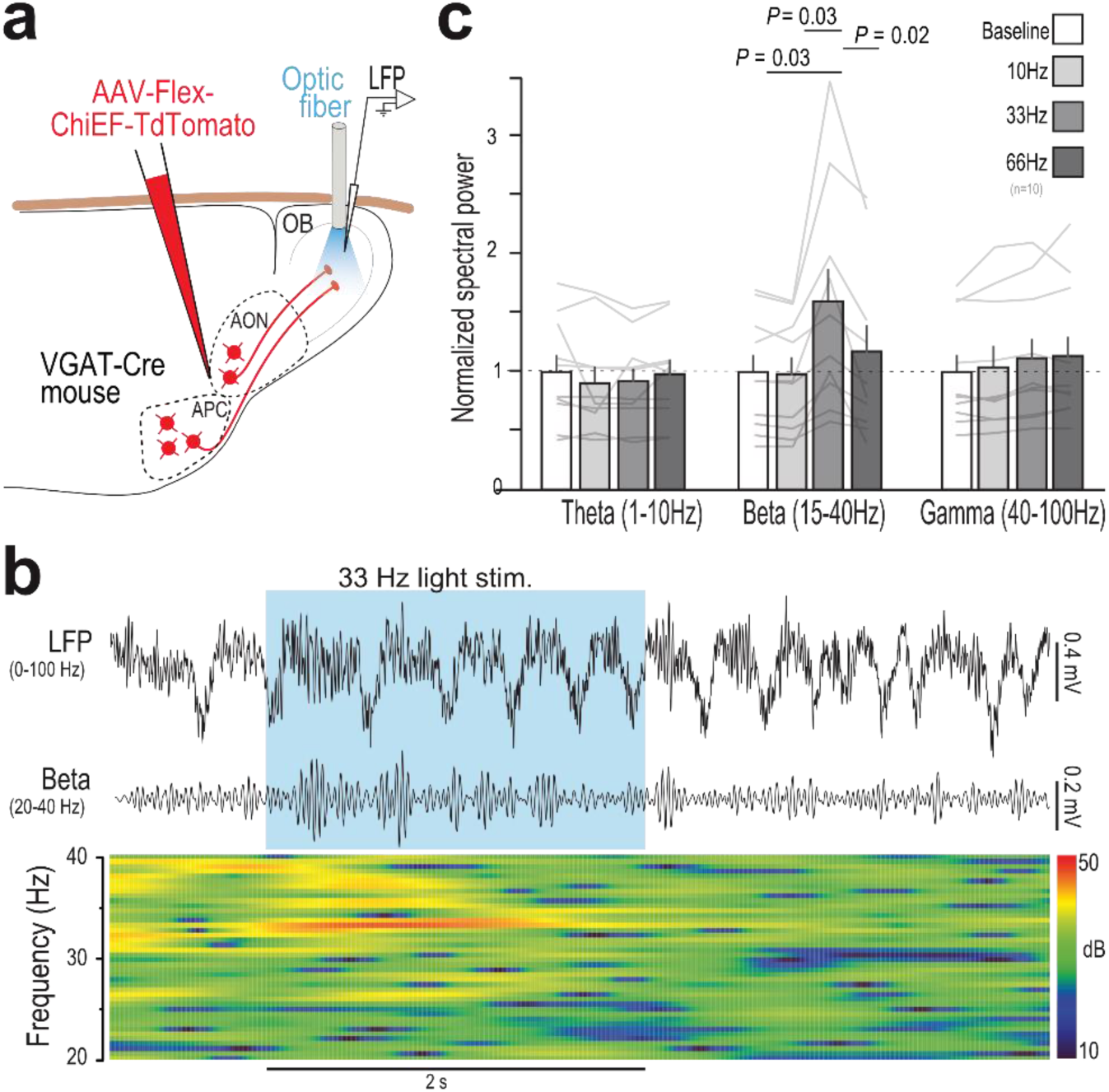
Cortico-bulbar GABAergic axon stimulation increases beta oscillations. **a**, Awake OB LFP recordings while GABAergic cortical axons were optogenetically stimulated through an optic fiber positioned in the dorsal OB in awake mice. **b**, Example broadband (1-100 Hz, up) and beta-filtered (20-40 Hz, middle) LFP trace recorded in the OB. Blue box: 33 Hz light stimulation (2 s). Bottom, corresponding time-frequency spectrogram of the LFP signal in the beta band. **c**, Quantification of the LFP band power (theta, beta and gamma) during 10 Hz, 33 Hz or 66 Hz stimulation patterns (n = 10 recording sites from 6 mice). Interaction between LFP band and stimulation patterns was significant (repeated measures two-way ANOVA, F(6,54)=8.08, *P*=10^-6^). Within the beta band, 33 Hz stimulation only had a significant effect (repeated measures one-way ANOVA, F(3,37)=9.875, p=0.0054; Tukey’s post-hoc test).

**Supplemental Fig. 7.**
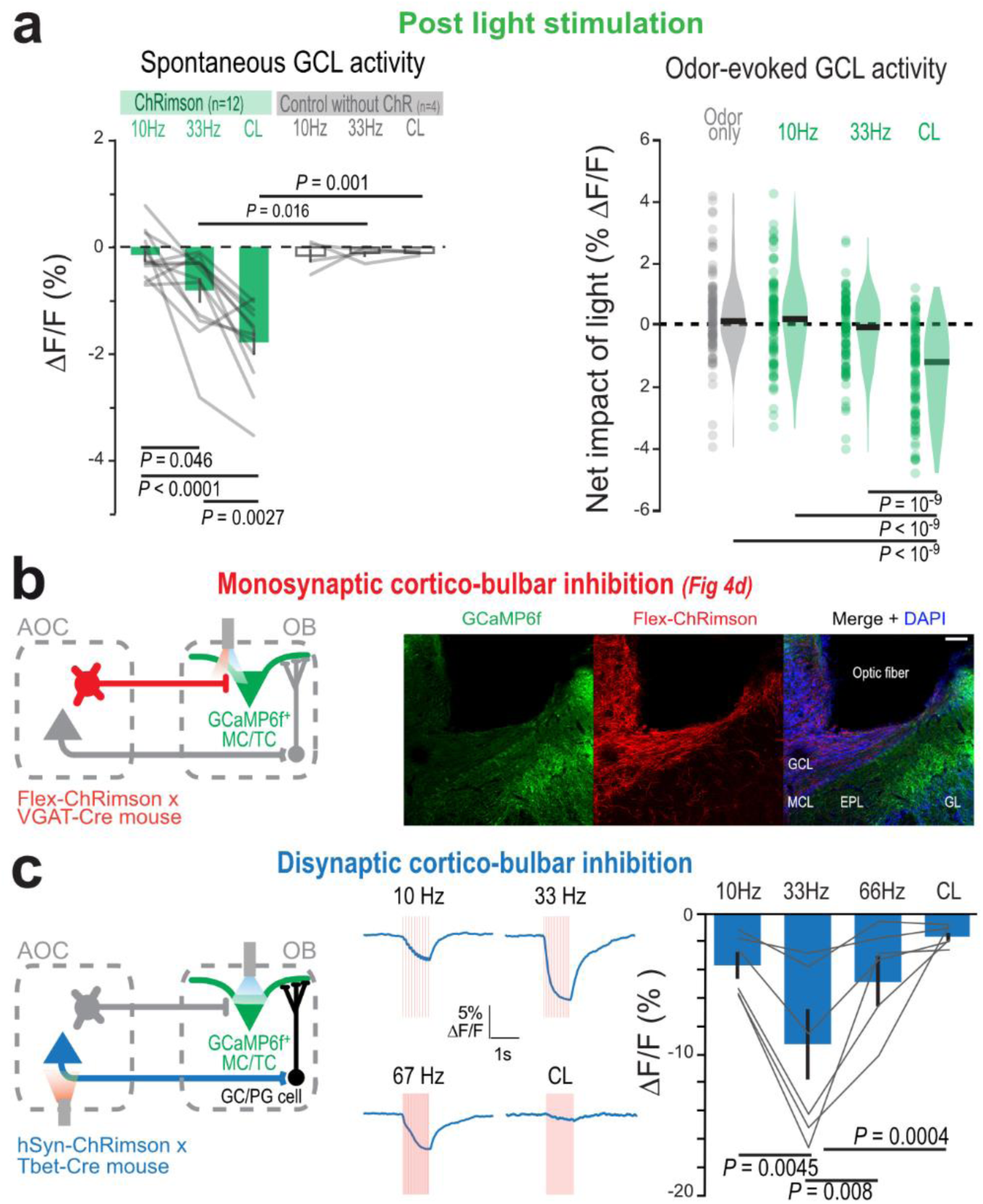
Further characterization of cortical GABAergic inhibition in the OB neuron populations. Inhibition induced by light-stimulation of ChRimson^+^ GABAergic AON/APC axons in the OB persisted 1 s after light stimulation offset. Left, spontaneous activity (RM-One-way-ANOVA with Tukey’s post-hoc test, F(2,22)=20.64, P<0.0001). Right, odor-evoked activity (One-way-ANOVA with Tukey’s post-hoc test). Data presented as mean ± sem; gray lines, individual mice. Violin plots are ks density estimates; black bar is median; circle, individual odor-recording site pair. **b**, Stimulation of GABAergic cortical feedback produces monosynaptic inhibition onto MC/TC populations (Fig. 4d). Right, Confocal image showing GCaMP6f expression in MCs and TCs, ChRimson-tdTomato in GABAergic cortical axons and the placement of the optic fiber above the lateral MCL and EPL. Blue, DAPI. Scale bar, 200 μm. **c**, Non-specific cortical feedback stimulation produces disynaptic inhibition onto MC/TC populations. Non-specific cortical feedback consists mainly in glutamatergic feedback (see Supplemental Fig.5). MC/TC population responses were collected utilizing fiber photometry in freely moving mice. In these experiments, we used Tbet-Cre mice to restrict GCaMP6f expression to MCs/TCs. **c**, Left, example averaged responses of MC/TC populations across different cortical feedback stimulation patterns. Right, U-shaped inhibition magnitude with increasing light stimulation frequency, up to CL. Cortical feedback stimulation at 33 Hz produced significantly more inhibition than any other stimulation pattern (RM-One-way ANOVA with Tukey’s post-hoc test, F(3,15)=10.89, P=0.0005, n = 6 recordings sites in 3 mice). This contrasted with the monotonic increase of inhibition magnitude with increasing stimulation frequency of cortical GABAergic feedback, up to CL. Data presented as mean ± sem; Gray, individual recording sites.

**Supplemental Fig. 8.**
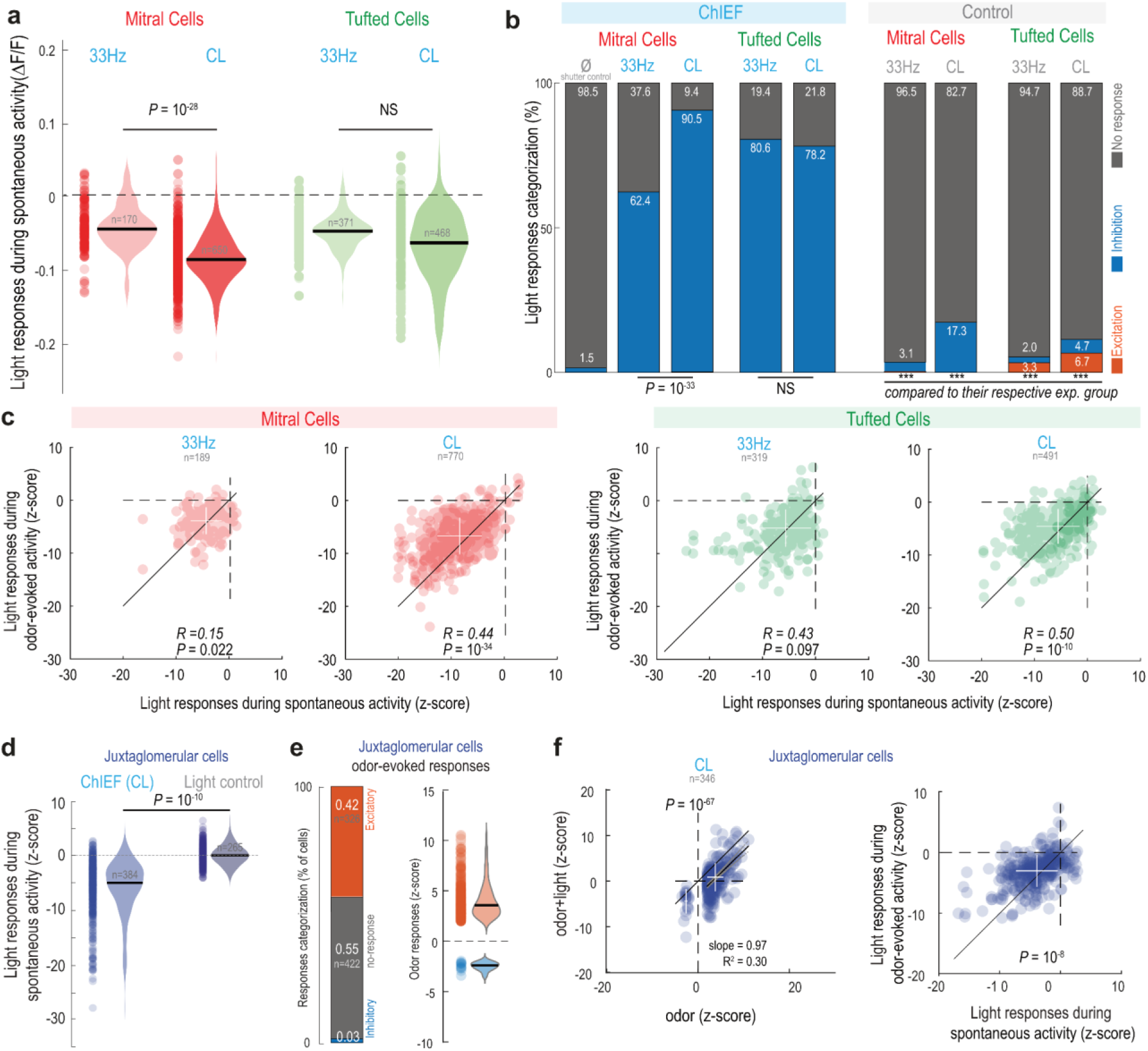
Additional analysis of cortico-bulbar inhibition of OB output neurons. **a**, ΔF/F0 light responses to 33 Hz and CL for MCs (left) and TCs (right) during spontaneous activity. All experimental groups were significantly higher than their respective light control (two-sided t-test, *P* << 0.001 for all comparisons). CL induced greater inhibition than 33 Hz stimulation in MCs, but not TCs (two-sided t-test). Violin represents ks-density estimation of the data. Black bar, median; circle, individual cell. n as in Fig. 5d. **b**, Categorization of light responses to optogenetic stimulation (left) and light control (right) for MCs and TCs during spontaneous activity. Light caused significantly more significant inhibition in all experimental groups compared to their respective light control (χ^2^ test, p << 0.001 for all comparisons). CL produced significant inhibition in a greater proportion of MCs, but not TCs (χ^2^ test). Note that control CL illumination on MCs induced signification inhibition in a sizable proportion of MCs. n as in Fig. 5d. **c**, Inhibition of spontaneous and odor-evoked activities were weakly correlated (Spearman’s correlation coefficient). In odor-responsive MCs (left) and TCs (right), light inhibition of spontaneous activity was slightly stronger than during odor-evoked (two-sided paired t-test), suggesting that the effect of cortical GABAergic feedback stimulation is context-dependent. White cross denotes mean ± s.d. n as in Fig. 5f. **d**, CL illumination induced a significant inhibition of JG spontaneous activity (z-scored) in the presence of ChIEF compared to the “light control” (One-way ANOVA with Tukey’s post-hoc test, *P* = 10^-10^). **e**, Categorization (left) and magnitude (right) of the odor-evoked responses in JG cells. Violins are ks-density estimates of the data. Black bar, median; circles, individual cell-odor pair. **f**, Effect of CL stimulation on odor-evoked responses in JG cells. Left: Odor-responses were significantly dampened upon CL stimulation (paired two-sided t-test). Right: Inhibition was slightly greater on spontaneous activity than during odor-evoked (two-sided paired t-test).

**Supplemental Fig. 9.**
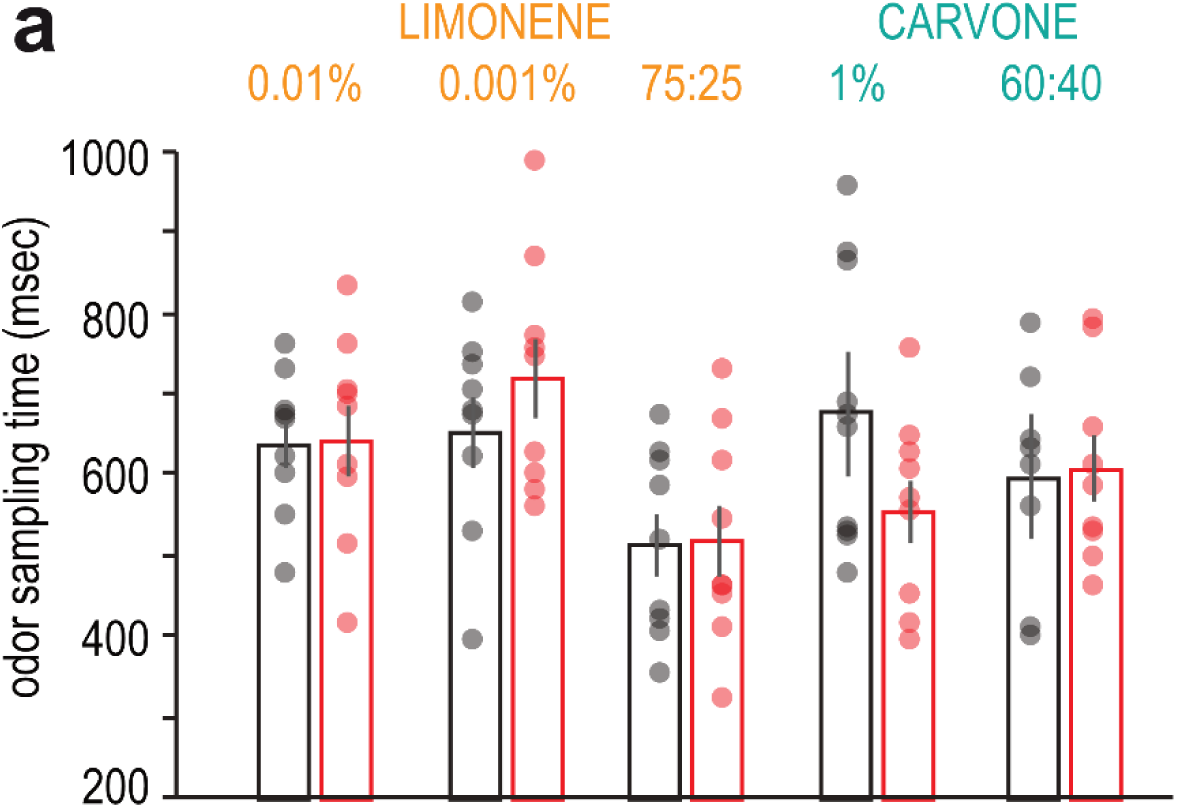
Additional analysis of the behavioral effects of GABAergic cortico-bulbar axon silencing. Mean odor sampling time for correct trials in sessions with mean performance on the last three blocks superior or equal to the criterion level (85%) for control (black) and DREADD groups (red).

**Supplemental Fig. 10.**
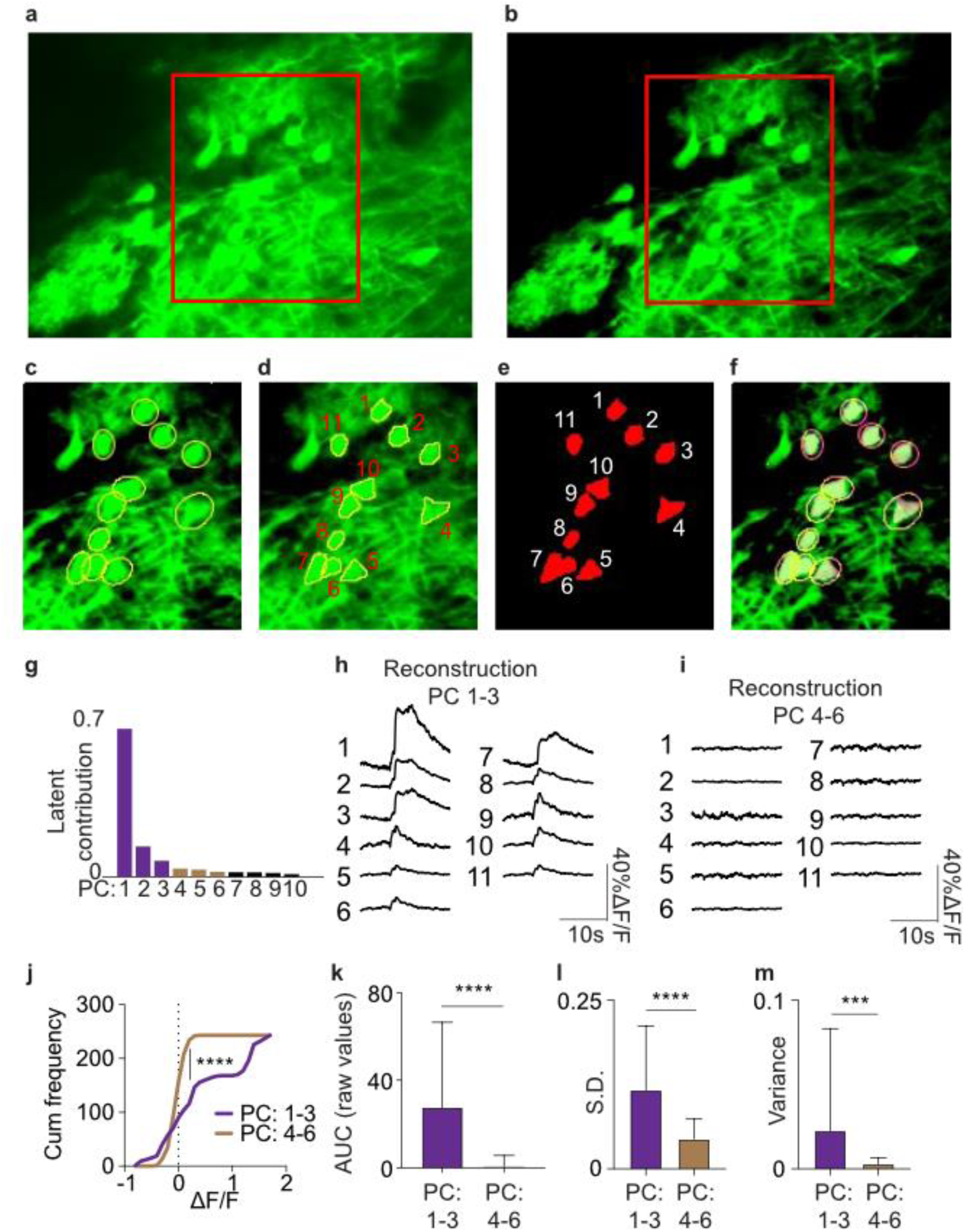
Protocol for neuronal activity data extraction using awake 2-photon imaging. **a**, maximum projection of a field of view from motion corrected raw images from a 2-photon recording session. **b,** PCA reconstruction of the same field of view in **a** to eliminate the background noise and allowing visualization for regions of interest (ROIs) selection. In **a** and **b**, the red boxes indicate the highlighted area in panels **c-f. c**, Manual selection of ROIs after the PCA-assisted reconstruction of the image series. **d,** ROI boundaries were redefined using a second PCA inside each ROI. For illustration purposes only, we show here the ROI reconstruction using the 1st PC only. Note that the new boundary only encompasses pixels smaller than the original elliptical ROIs. **e,** The ROI shape after the reconstruction using PCA inside the elliptical ROIs. **f,** Superimposition of the original manually drawn elliptical ROIs and the PCA-corrected ROIs. **g,** Eigenvalue contribution plot indicating that the first 3 PCs inside the ROI capture most of the variation in the dataset. PC1-3 were used to define the ROI boundaries in our study. Traces from the PCA-corrected ROIs in **d** using either PCs 1-3, capturing the most significant variability in the original dataset (**h**) or PCs 4-6, capturing the ‘noisy’ contributions in the dataset (**i**). Traces are from the ROIs in **e. j,** Cumulative frequency of the ΔF/F values using PCs 1-3 (purple) or PCs 4-6 (yellow). Note that the ΔF/F values from PCs 4-6 reconstruction are close to 0 and are significantly different from the ΔF/F value distribution from PCs 1-3 reconstruction (p < 0.0001). Area under the (AUC) as a measure of activity (**k**), standard deviation (**l**) and variance (**m**) of the ΔF/F values for the reconstructed data from PCs 1-3 are significantly higher than those contributing to the noisy pixels, PCs 4-6 (***: p < 0.0001; **: p < 0.001).

**Supplemental Table 1.** Membrane resistance (Rm) and kinetics of the light-evoked IPSCs in the post-synaptic neurons. Data presented as mean ± sem.

|  | **Rm (MOhm)** | **Latencies (ms)** | **Rise time (ms)** | **Tau (ms)** |
| --- | --- | --- | --- | --- |
| **GC** | 875.90 ± 117.30 (n = 33) | 3.04 ± 0.27 (n = 31) | 0.40 ± 0.04 (n = 17) | 9.64 ± 1.85 (n = 15) |
| **dSAC** | 268.60 ± 55.71 (n = 10) | 2.88 ± 0.31 (n = 8) | 0.43 ± 0.09 (n = 7) | 10.35 ± 1.77 (n = 6) |
| **MC** | 72.96 ± 7.26 (n = 19) | 4.26 ± 0.42 (n = 13) | 1.01 ± 0.16 (n = 10) | 9.20 ± 1.60 (n = 6) |
| **TC** | 73.27 ± 10.31 (n = 11) | 4.74 ± 0.18 (n = 11) | 0.87 ± 0.11 (n = 12) | 10.11 ± 2.36 (n = 9) |
| **eTC** | 149.30 ± 29.18 (n = 16) | 3.33 ± 0.40 (n = 9) | 0.70 ± 0.22 (n = 8) | 5.88 ± 2.71 (n = 5) |
| **PG cell** | 584.20 ± 46.71 (n = 29) | n/a | n/a | n/a |
| **sSAC** | 264.60 ± 50.78 (n = 9) | n/a | n/a | n/a |

